# Single cell transcriptomics reveals correct developmental dynamics and high-quality midbrain cell types by improved hESC differentiation

**DOI:** 10.1101/2022.09.15.507987

**Authors:** Kaneyasu Nishimura, Shanzheng Yang, Ka Wai Lee, Emilía Sif Ásgrímsdóttir, Kasra Nikouei, Wojciech Paslawski, Sabine Gnodde, Guochang Lyu, Lijuan Hu, Carmen Saltó, Per Svenningsson, Jens Hjerling-Leffler, Sten Linnarsson, Ernest Arenas

**Affiliations:** Department of Medical Biochemistry and Biophysics, Karolinska Institutet, Stockholm 171 77, Sweden; Department of Clinical Neuroscience, Karolinska University Hospital, Stockholm 171 77, Sweden

**Author notes:** Laboratory of Functional Brain Circuit Construction, Graduate School of Brain Science, Doshisha University, Kyoto, 610-0394, Japan (Kaneyasu Nishimura). These authors equally contributed to this work (Shanzheng Yang, Ka Wai Lee). Corresponding author: Ernest Arenas, M.D., Ph.D., Department of Medical Biochemistry and Biophysics, Karolinska Institutet, 17177 Stockholm, Sweden.

**Keywords:** Human midbrain development, human embryonic stem cell, midbrain dopaminergic neuron, single-cell RNA sequencing, Parkinson’s disease

## Abstract

Stem cell technologies provide new opportunities for modeling cells in the healthy and diseased states and for regenerative medicine. In both cases developmental knowledge as well as the quality and molecular properties of the cells are essential for their future application. In this study we identify developmental factors important for the differentiation of human embryonic stem cells (hESCs) into midbrain dopaminergic (mDA) neurons. We found that Laminin-511, and dual canonical and non-canonical WNT activation followed by GSK3β inhibition plus FGF8b, improved midbrain patterning. In addition, mDA neurogenesis and differentiation was enhanced by activation of liver X receptors and inhibition of fibroblast growth factor signaling. Moreover, single-cell RNA-sequencing analysis revealed a developmental dynamics similar to that of the endogenous human ventral midbrain and the emergence of high quality molecularly-defined midbrain cell types, including mDA neurons that become functional. Thus, our study identifies novel factors important for human midbrain development and opens the door for a future application of molecularly-defined hESC-derived midbrain cell types in Parkinson’s disease.

## Introduction

Midbrain dopaminergic (mDA) neurons are known to control several important functions in humans, such as voluntary movement, cognition, motivation and reward. Amongst them, mDA neurons of the substantia nigra pars compacta (SNc) project to the caudate-putamen and form the nigrostriatal pathway, which controls voluntary movements. The loss of SNc DA neurons and of dopamine in the caudate-putamen is a defining feature of Parkinson’s disease (PD) (Damier et al., 1999), a neurodegenerative disorder characterized by paucity of movements, tremor, rigidity and loss of postural control (Lees et al., 2009). However, the cause of PD is largely unknown and current treatments are symptomatic and loose efficiency with time.

Progress in understanding the molecular logic and mechanisms controlling mDA neuron development has led to important developments in different areas of stem cell biology, including PD modeling, drug screening and personalized therapeutics (Caiazza et al., 2020), as well as PD cell replacement therapy (Adler et al., 2019; Arenas et al., 2015; Doi et al., 2020; Kikuchi et al., 2017; Kim et al., 2021; Kirkeby et al., 2017; Moriarty et al., 2022; Schweitzer et al., 2020; Tao et al., 2021). mDA neurons are currently thought to derive from radial glia-like progenitor cells at the caudal and ventral end of the midbrain floorplate (Bonilla et al., 2008; Ono et al., 2007). This area is controlled by signals derived from two organizing centers, the midbrain-hindbrain boundary (MHB) and the floor plate (Wurst et al., 2001). One of the most critical signaling events in mDA neuron development is the activation of the Wnt/μ-catenin pathway by Wnt1, a morphogen derived from these two centers (Arenas, 2014). Wnt1 controls several aspects of mDA neuron development such as anterior-posterior patterning (McMahon and Bradley, 1990; Thomas and Capecchi, 1990), the specification of mDA progenitors (Prakash et al., 2006) and the induction of mDA neurogenesis in the midbrain floorplate (mFP) (Andersson et al., 2013). Accordingly, activation of this pathway in human pluripotent stem cells (hPSCs) by glycogen synthase kinase (GSK)3μ inhibitors, such as CHIR99021, has led to significant improvements in protocols for the generation of mDA neurons (Denham et al., 2012; Doi et al., 2014; Kim et al., 2021; Kriks et al., 2011). However, there are multiple additional developmental factors and signaling pathways known to control mDA neuron development in mice, whose function in human midbrain development remains to be examined. One of them is Wnt5a, a morphogen known to promote midbrain morphogenesis, neurogenesis and mDA progenitor differentiation in the developing mouse midbrain (Andersson et al., 2008; Castelo-Branco et al., 2003). Wnt5a is known to activate the Wnt/planar cell polarity/Rac1 (Wnt/PCP/Rac1) pathway in mDA progenitors and neurons (Andersson et al., 2008; Čajánek et al., 2013; Parish et al., 2008). Moreover, analysis of double *Wnt1* and *Wnt5a* knockout mice revealed a complex interplay between these two pathways, which controls diverse aspects of ventral midbrain development (Andersson et al., 2013; Arenas, 2014). Another interesting pathway that remains to be implemented in advanced protocols for mDA differentiation of human embryonic stem cells (hESCs) is activation of the nuclear receptors NR1H3 and NR1H2 (also known as liver X receptor α and β, LXRs), which control not only lipid metabolism but also mDA neurogenesis both *in vitro* and *in vivo* (Sacchetti et al., 2009; Theofilopoulos et al., 2013; Toledo et al., 2020). Finally, one additional component that we examined is the midbrain-specific extracellular matrix protein, laminin 511 (LN511), which is known to expand hPSC-derived mDA progenitors (Doi et al., 2014; Kirkeby et al., 2017) and differentiate neuroepithelial stem cells (Zhang et al., 2017), but it is unclear whether full length LN511 can control progenitor identity and mDA differentiation in hESCs.

Single-cell RNA-sequencing (scRNA-seq) has provided very powerful and unbiased insights into the cell types and gene expression profiles in the developing human ventral midbrain (Andersson et al., 2006; Birtele et al., 2020; La Manno et al., 2016). In our study, we therefore used scRNA-seq data (La Manno et al., 2016) as a blueprint of the developmental dynamics and the cell types physiologically found in the developing midbrain *in vivo*, as well as a reference dataset to evaluate cell composition and quality of cell types generated by hESCs during mDA differentiation. Four different types of endogenous human progenitors have been found in the endogenous human midbrain floorplate: The ventral midline progenitor (ProgM), medial floorplate progenitor (ProgFPM), lateral floorplate progenitor (ProgFPL), and neuronal progenitor (NProg). In addition, two radial glia-like cells (Rgl1 and Rgl3) were found enriched in the ventral midbrain floorplate. Moreover, four of these cell types (ProgM, ProgFPM, ProgFPL and Rgl1) are known to selectively express key factors required for the specification of mDA progenitors: the morphogen *WNT1,* and the transcription factors *LMX1A*, *OTX2* and *FOXA2,* all of which are required for mDA neuron development (Andersson et al., 2006; Ferri et al., 2007; Puelles et al., 2004). Other cells of interest are the neuronal progenitor (NProg) and the first postmitotic cell of the mDA lineage, the medial neuroblast (NbM), both of which express genes either involved in or required for mDA neurogenesis, such as *NEUROD1* and *NEUROG2* (*NGN2*) (Kele et al., 2006), respectively. The NbM also expresses the nuclear receptor *NR4A2* (*NURR1*), which is required for mDA neuron development (Zetterström et al., 1997). *NR4A2* is also expressed in the three embryonic mDA neuron subpopulations (DA0, DA1 and DA2) (La Manno et al., 2016), together with tyrosine hydroxylase (*TH*) and transcription factors required for mDA development, such as Engrailed 1 (*EN1*) (Simon et al., 2001), Pre-B-cell leukemia homeobox 1 (*PBX1*) (Villaescusa et al., 2016), and Pituitary homeobox 3 (*PITX3*) (Nunes et al., 2003). However, despite all this knowledge, the precise cell composition and quality of hESC-derived midbrain cell types compared with endogenous single cell standards remains largely undefined.

In this study we leverage existing human scRNA-seq data and functional analysis of the developing mouse midbrain, to explore whether three key developmental components (WNT5A, LXR and LN511) can improve the generation of mDA neurons from hESCs. We carefully monitor pathway activation by synchronizing gene expression in hESCs differentiating into mDA neurons with that in endogenous human ventral midbrain development. We found that dual activation of Wnt/μ-catenin and Wnt/PCP/Rac1 with CHIR99021 and WNT5A, respectively, together with activation of LXRs with a synthetic ligand and extended use of LN511 improves mDA differentiation of hESCs. Moreover, scRNA-seq allowed us to define the quality of the hESC-derived cells compared with the endogenous human ventral midbrain standards. We found that our human development-based protocol recapitulates key features of human midbrain development, including the generation ventral midbrain cell types similar to those in the developing human ventral midbrain, and functional mDA neurons. Our study thus defines the function of developmental factors during human midbrain differentiation and shows their implementation improves the composition and quality of hESC-derived mDA cultures.

## Results

### Efficient induction of midbrain floor plate progenitors by LN511 and dual WNT activation

Human ESCs were cultivated in chemically-defined medium with the dual Smad inhibitors LDN193189 and SB431542 to promote neural induction (Chambers et al., 2009), the Shh agonist purmorphamine to ventralize (Kriks et al., 2011), and the GSK3β inhibitor CHIR99021 to activate Wnt/βcatenin signaling and achieve caudal midbrain identity (Figure 1A) (Imaizumi et al., 2015; Kim et al., 2021; Kirkeby et al., 2012; Kriks et al., 2011). ScRNA-seq data of the developing human ventral midbrain was used to monitor both midbrain patterning and the emergence of midbrain progenitor markers (*LMX1A, FOXA2* and *OTX2)*, which are expressed by four different cell types, Rgl1, ProgM, ProgFPL and ProgFPM (Figure 1B) (La Manno et al., 2016). Quantitative PCR (qPCR) analysis at day 11 (Figure 1C and **Figure S1A**) showed that 2.0-3.5 µM CHIR99021 induced the expression of hindbrain marker genes (*FGF8B*, *GBX2* and *HOXA2*), while ventral midbrain genes (*LMX1A, FOXA2, OTX2* and *CORIN*) were upregulated by 1.0-1.5 µM CHIR99021. Moreover, 1.5 µM CHIR99021 increased the expression of caudal ventral midbrain genes (*MSX1, WNT5A, WNT1* and *EN1*). Since dopaminergic progenitors are enriched in the caudal midbrain (Kirkeby et al., 2017), we chose 1.5 µM CHIR99021 as a baseline to examine the possible function of factors whose function in early mDA progenitor patterning has not yet been fully established, such as LN511 and WNT5A.

**Figure 1.**
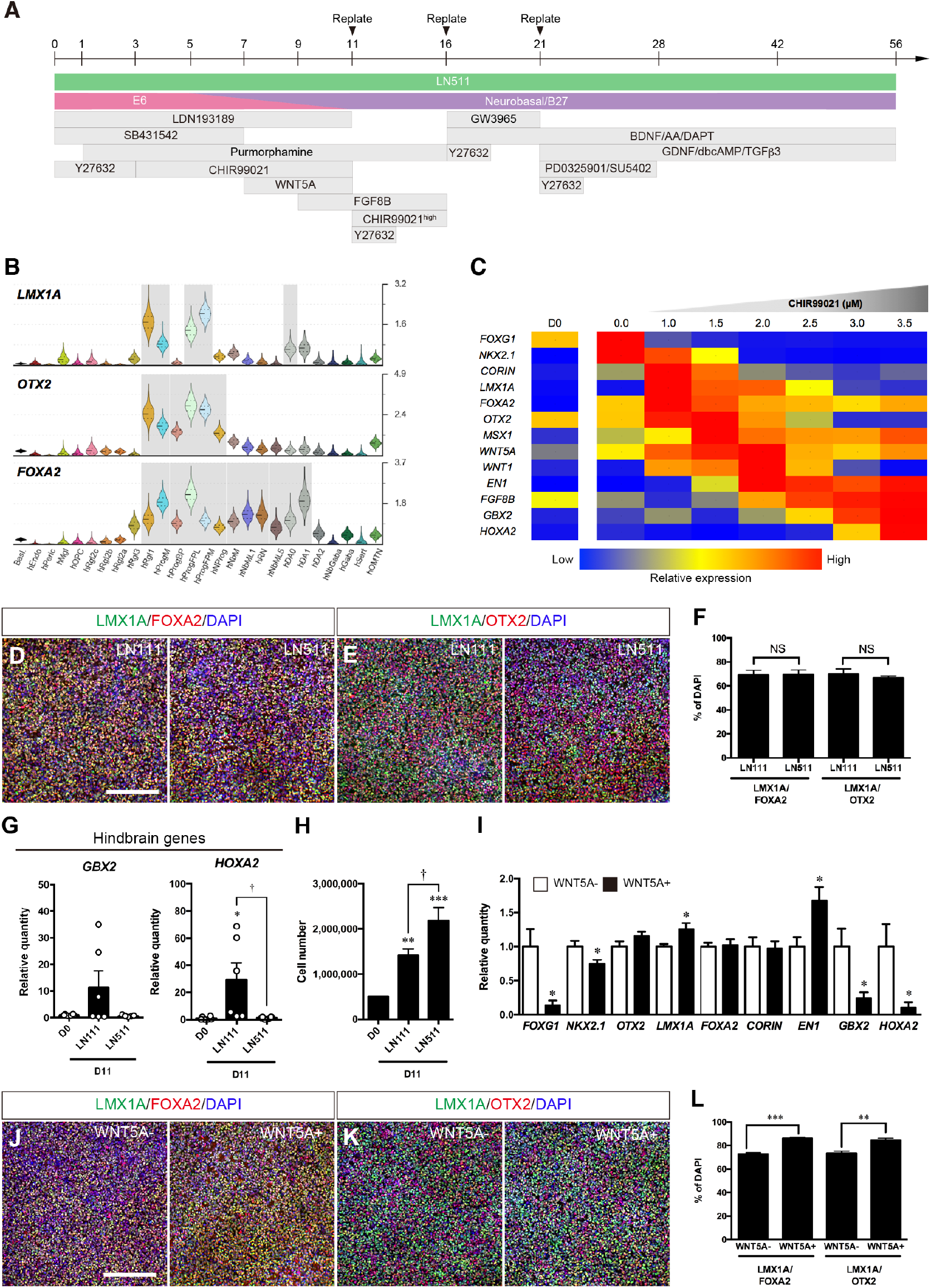
Induction of floorplate progenitors from hESCs at day 11 using a new developmental-based protocol. (**A**) Schematic of the differentiation protocol for midbrain DA neurons. (**B**) Violin plots LMX1A, FOXA2 and OTX2 generated from scRNA-seq data of developing human ventral midbrain shown across corresponding cell types. Right axis shows absolute molecular counts. Grey, enriched over baseline with posterior probability >99.8%. For cell type nomenclature, see La Manno et al., (2016). (**C**) Gene expression profile of differentiated cells according to CHIR99021 concentration at day 11. (**D** and **E**) Immunostaining of LMX1A^+^/FOXA2^+^ cells (**D**) and LMX1A^+^/OTX2^+^ cells (**E**) at day 11. Scale bar, 200 µm. (**F**) Quantification of LMX1A^+^/FOXA2^+^ cells and LMX1A^+^/OTX2^+^ cells. NS, not significant (n = 4). (**F**) qPCR analysis of *GBX2* and *HOXA2* of differentiating cells on either LN111 or LN511. *p < 0.05 vs. D0; †p < 0.05 vs LN511 (n = 6). (**H**) The cell yield of differentiating cells on either LN111 or LN511 on day 11. **p < 0.01, ***p < 0.001 vs. D0; †p < 0.05 vs LN511 (n = 6). (**I**) qPCR analysis of the differentiated cells presence/absence of WNT5A at day11 (n = 5-9). *p < 0.05 vs. WNT5A(-) condition. (**J** and **K**) Immunostaining of LMX1A^+^/FOXA2^+^ cells (**J**) and LMX1A^+^/OTX2^+^ cells (**K**) at day 11. Scale bar, 200 µm. (**L**) Quantification of LMX1A^+^/FOXA2^+^ cells and LMX1A^+^/OTX2^+^ cells. ***p < 0.001 vs. WNT5A(-) condition (n = 4).

We first focused on the extracellular matrix protein LN511, which is enriched in the developing human ventral midbrain and is known to promote the differentiation and survival of mDA neurons (Zhang et al., 2017). hESCs were cultivated on good manufacturing practice (GMP)-grade LN111 or LN511, until day 11. In both cases ≈70% of the cells were LMX1A^+^/FOXA2^+^ cells and LMX1A^+^/OTX2^+^ (Figure 1D-1F) and the pluripotent stem cell markers *NANOG* and *POU5F1* drastically decreased (Figure 1B). However, LN511 but not LN111 significantly decreased the expression of hindbrain markers such as *GBX2* and *HOXA2* (Figure 1G). In addition, a greater yield of mDA progenitor cells was obtained with LN511 than on LN111 (Figure 1H). Thus our results show that LN511 efficiently expands midbrain progenitors and prevents the expression of hindbrain patterning genes.

We next investigated the function of WNT5A, a morphogen co-expressed with WNT1 in the four candidate human ventral midbrain DA progenitors (Rgl1, ProgM, ProgFPM and ProgFPL) (La Manno et al., 2016). Since Wnt1 and Wnt5a are known to cooperate to promote mDA neuron development *in vitro* and *in vivo* (Andersson et al., 2013; Castelo-Branco et al., 2003), we performed a dual WNT activation of hESCs with CHIR99021 and WNT5A from day 7 to day 11 and then examined patterning markers. qPCR analysis at day 11 revealed a significant decrease in the expression of *FOXG1*, *NKX2.1*, *GBX2* and *HOXA2* and a significant increase in *LMX1A* and *EN1* after treatment with WNT5A (100 ng/mL) (Figure 1I). Moreover, WNT5A also increased the proportion of LMX1A^+^/FOXA2^+^ and LMX1A^+^/OTX2^+^ cells in a significant manner, to reach 86.1 ± 0.7% and 84.6 ± 1.7%, respectively (Figure 1J-1L). This regulation was specific as it did not change the proportion of CORIN^+^ or LMX1A^+^/CORIN^+^ cells (Figure 1C and 1D), two markers expressed in ProgM (La Manno et al., 2016). These results indicate that WNT5A, in combination with CHIR99021 (1.5 µM), improves the induction of midbrain DA progenitors compared with CHIR99021 alone at the expense of more anterior and posterior fates. Moreover, comparable midbrain patterning was confirmed in three different hESC lines, HS401, HS975 and HS980 (Figure 1E-1H), indicating that the effects of dual WNT activation are both specific and robust.

### Specification of caudal-ventral midbrain domain by CHIR99021 and fibroblast growth factor 8b (FGF8b)

It is known that mDA neurons originate in the caudal floorplate domain under the influence of Wnt1, a morphogen strongly expressed in the midbrain side of the midbrain-hindbrain boundary (MHB) and in the two bands that define the lateral floorplate (Burbach and Smidt, 2006; Prakash et al., 2006; Wurst et al., 2001). In addition the hindbrain side of the MHB expresses FGF8b, a factor also known to induce mDA neurons (Ye et al., 1998). We therefore treated our cultures with both CHIR99021 and FGF8b in the presence of purmorphamine, to mimic the morphogens controlling the midbrain floorplate. To estimate the resulting strength of Wnt signaling during CHIR99021, purmorphamine and FGF8b treatment, we examined the expression of endogenous canonical and non-canonical Wnts (Figure 2A). As expected, activation of canonical Wnt/β-catenin signaling with increasing concentrations of CHIR99021 downregulated the expression of *WNT1* and *WNT7A*, but did not affect the expression of *WNT5A* and *WNT11*. Notably, 7.5 µM CHIR99021 did not change the proportion of LMX1A^+^/FOXA2^+^ cells and LMX1A^+^/OTX2^+^ cells (Figure 2A-2C), but increased the expression of *EN1* (Figure 2B) and EN1-immunoreactivity in LMX1A^+^ cells compared with 1.5 µM CHIR99021 (Figure 2C and 2D). Moreover rostral and/or lateral midbrain markers such as *NKX2.1, BARHL1, PITX2* and *SIX3* decreased with the concentration of CHIR99021 (Figure 2E). Thus our results indicate that the combination of 7.5 µM CHIR99021 and FGF8b from day 11 to day 16 does not only effectively increase caudal midbrain gene expression, but also decreases rostral and lateral gene expression.

**Figure 2.**
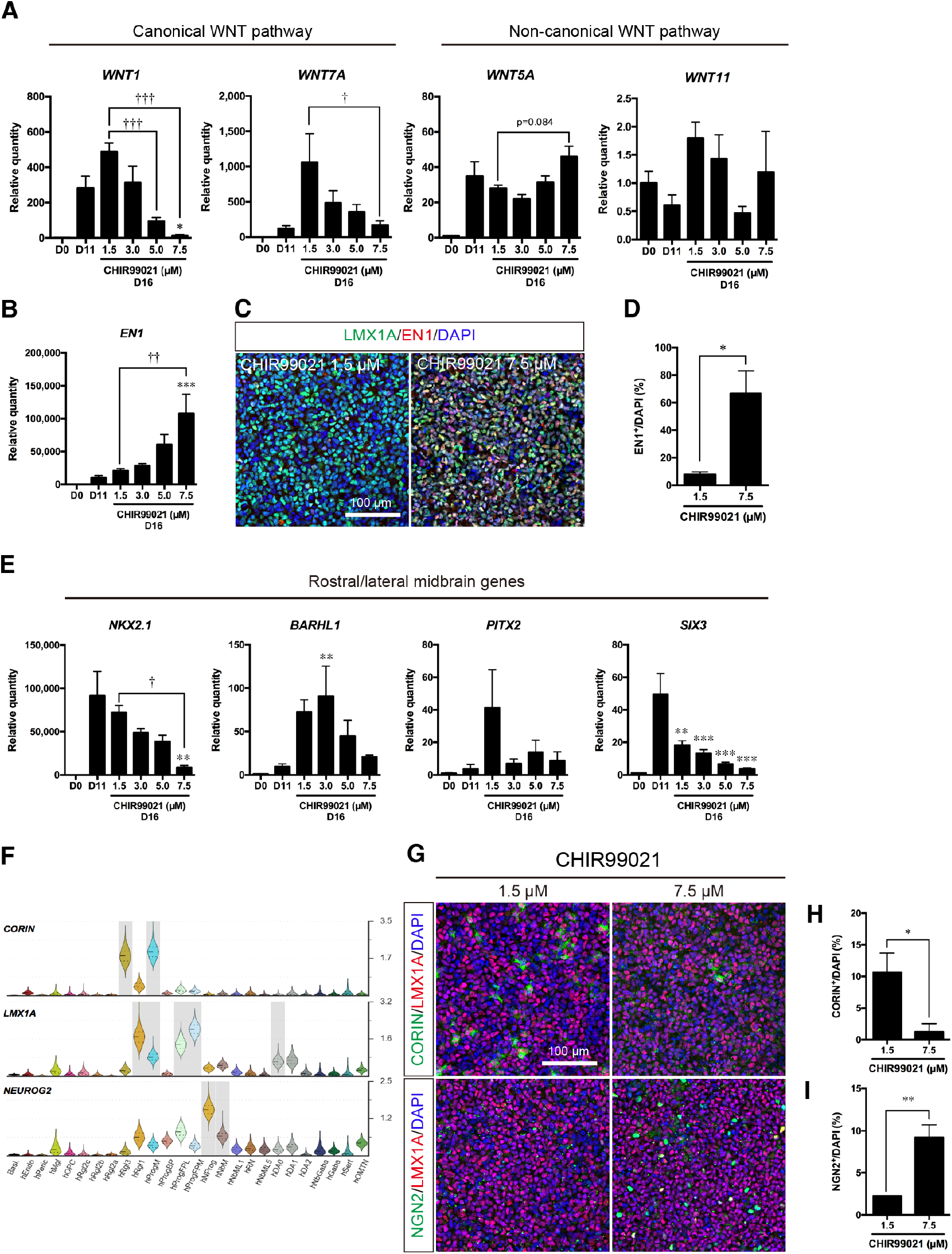
Analysis of floor plate patterning of hESC-derived neural progenitors at day 16. (**A**) qPCR analysis of canonical WNT pathway and non-canonical WNT pathway at day 16. *p < 0.05 vs. D11; ††p < 0.01, †††p < 0.001 vs. 1.5 µM CHIR99021 (n = 6). (**B**) qPCR analysis of EN1 at day 16. ***p < 0.001 vs. D11; ††p < 0.01 vs. 1.5 µM CHIR99021 (n = 6). (**C**) Immunostaining of LMX1A^+^/EN1^+^ cells. Scale bar, 100 µm. (**D**) Quantification of LMX1A^+^/EN1^+^ cells. *p < 0.05 vs. 1.5 µM CHIR99021 (n = 3). (**E**) qPCR analysis of rostral and lateral midbrain markers at day 16. **p < 0.01, ***p < 0.001 vs. D11; †p < 0.05 vs. 1.5 µM CHIR99021 (n = 6). (**F**) Violin plots of CORIN, LMX1A and NGN2 generated from scRNA-seq data of developing human ventral midbrain. (**G**) Immunostaining of CORIN^+^/LMX1A^+^ cells and NGN2^+^/LMX1A^+^ cells. Scale bar, 100 µm. (**H** and **I**) Quantification of CORIN^+^/LMX1A^+^ cells (**H**) and NGN2^+^/LMX1A^+^ cells (**I**). *p < 0.05, **p < 0.01 vs. 1.5 µM CHIR99021 (n = 3).

We also investigated whether combined 7.5 µM CHIR99021 and FGF8b promotes lineage progression from proliferative to neurogenic progenitors. We therefore examined the presence of cells expressing *CORIN*, an early midbrain gene selectively expressed in Rgl3 and ProgM cell types, and *NGN2* a gene selectively expressed in two cell types undergoing neurogenesis, NProg and NbM (Figure 2F). Analysis of hESCs treated with 7.5 µM CHIR99021 day 11-16 revealed a decrease in CORIN^+^ cells, and an increase in NGN2^+^ cells, compared to 1.5 µM CHIR99021 (Figure 2G-2I). In addition, we also observed a very low proportion of LMX1A^+^/CORIN^+^ ProgM cells and a higher proportion of LMX1A^+^/NGN2^+^ cells in three different hESC lines treated with 7.5 µM CHIR99021 (Figure 2D-2H). Thus, combined, our results indicate that treatment with 7.5 µM CHIR99021 and FGF8 promotes cell lineage progression towards neurogenesis.

### Promotion of neurogenesis in hESC-derived progenitors by LXR activation

On day 16, our cultures contained abundant LMX1A^+^, FOXA2^+^ and CORIN^-^ cells, indicating that most of the cells are ProgFPL and ProgFPM. On the other hand, we observed a growing number of NGN2^+^ cells within the LMX1A^+^/FOXA2^+^ population, suggestive of an emerging NProg population. We thus examined whether neurogenesis could be enhanced by the synthetic LXR ligand, GW3965. Treatment with GW3965 (5-10 μM, day16-21) upregulated the expression of the LXR target genes, *SREBF1* (Figure 3A) and *ABCA1* (Figure 3B), indicating effective activation of LXRs. In addition we monitored the expression of *SOX2,* a neural progenitor marker, and of doublecortin (*DCX*), a marker expressed during neurogenesis in NProg and in all postmitotic neuroblasts and neurons (Figure 3C). We found that 10 µM GW3965 significantly decreased the proportion of SOX2^+^ progenitors, and increased the proportion of DCX^+^ cells (Figure 3D-3G). Moreover, 5-ethynyl-2’-deoxyuridine (EdU) pulse-chase experiments revealed an increase in neurogenesis as shown by the increased proportion of EdU^+^/DCX^+^ cells by 10 µM GW3965 at day 21 (Figure 3G and 3H). However, at this stage, immature cells including progenitors and NGN2^+^ were still present in the cultures (Figure 3I) and TH^+^ neurons had not yet emerged.

**Figure 3.**
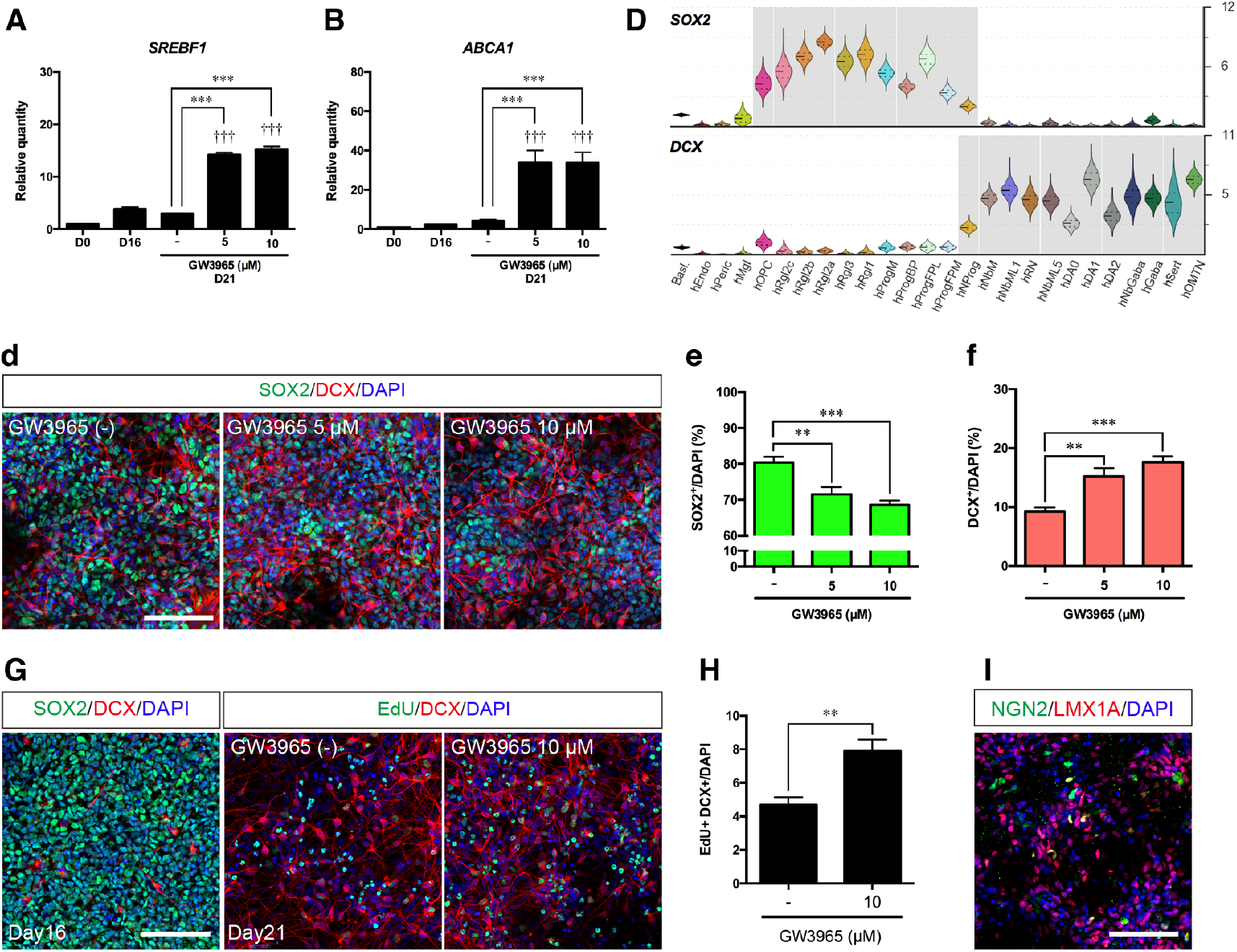
Analysis of hESC differentiation and neurogenesis at day 21. qPCR analysis of *SREBF1* (**A**) and *ABCA1* (**B**) at day 21. ***p < 0.001 vs. GW3965(-) condition. †††p < 0.001 vs. day 16 (n = 6). (**C**) Violin plots of SOX2 and DCX generated from scRNA-seq data of developing human ventral midbrain. (**D**) Immunostaining of SOX2^+^/DCX^+^ cells at day 21. Scale bar, 100 µm. (**E** and **F**) Quantification of SOX2^+^ cells (**E**) and DCX^+^ cells (**F**). **p < 0.01 vs. GW3965 (-) condition (n = 8). (**G**) Immunostaining of SOX2^+^/DCX^+^ cells at day 16 and EdU^+^/DCX^+^ cells at day 21, after GW3956 treatment (day 16-21). EdU pulse was performed for 4 hr at day 16 and EdU detection was performed at day 21. Scale bar, 100 µm. (**H**) Quantification of EdU^+^/DCX^+^ cells at day 21. **p < 0.01 vs. GW3965(-) condition (n = 6). **(I**) Immunostaining of LMX1A^+^/NGN2^+^ cells at day 21. Scale bar, 100 µm.

### Maturation of hESC-derived neurons by blocking of FGF signaling

FGF receptors 1-3 are predominantly expressed in immature *SOX2*^+^ cell types in the developing human ventral midbrain, such as radial glia and progenitors (Figure 4A). Since FGF signaling is important to maintain and expand neural precursors (Elkabetz et al., 2008; Koch et al., 2009), we speculated that inhibition of FGF signaling may limit the growth of progenitors and promote their differentiation. To inhibit FGF signaling, cultures were treated from day 21 to day 28 with 1 µM PD0325901, a MEK/ERK pathway inhibitor, and 5 µM SU5402, an FGF receptor inhibitor. We found that treatment with PD0325901 and SU5402 drastically reduced SOX2^+^ cell clusters and the number of phospho-histone H3 (pH3)^+^ cells at day28 (Figure 4B). At this stage, some cells exhibited neuronal morphology and expressed TH together with either LMX1A or FOXA2, suggesting the emergence of the dopaminergic DA0 population (Figure 4C). Markers identified at the single cell level in SNc neurons and in embryonic dopaminergic neurons type 2 (DA2) (La Manno et al., 2016), such as *LMO3* and *ALDH1A1* (Figure 4D), were examined by qPCR and were found significantly increased at day 28 after treatment with PD0325901 and SU5402 (Figure 4E and 4F). These results indicate that blocking FGF signaling promotes the maturation of the cultures and the emergence of mDA neurons. In agreement with these findings, a time-course analysis of gene expression by qPCR confirmed that the expression of progenitor markers (*SOX2*, *NEUROG2*, *LMX1A* and *FOXA2*) peaked at day 21 and decreased at day 28, while markers of postmitotic cells (*DCX* and *TUBB3*) and of mDA neurons (*NR4A2* and *POU6F1*) peaked at day 28 and remained stable thereafter (Figure 3A and 3B). Moreover, markers and transcription factors expressed at the single cell level in mDA neurons (*TH, KNCJ6, CALB1, ERBB4* and *PBX1*) or selectively in DA2 neurons (such as *LMO3, DEAF1* and *DKK3*) (La Manno et al., 2016), increased at days 42 and 56, suggesting a stable generation of postmitotic mDA neurons (Figure 4G).

**Figure 4.**
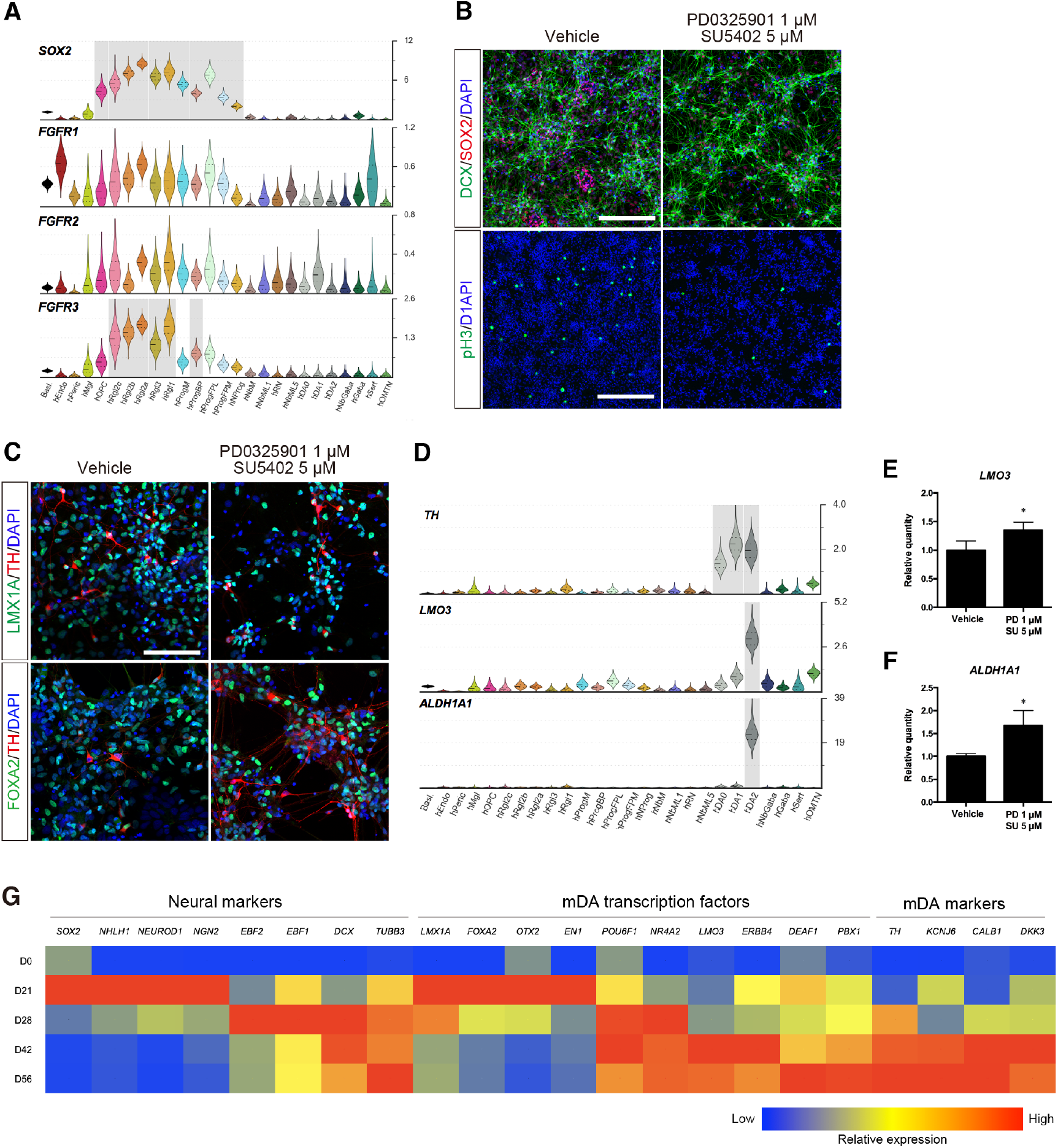
Analysis of postmitotic cells at day 28 and mDA neuron differentiation. (**A**) Violin plots of SOX2 and FGFRs generated from scRNA-seq data of developing human ventral midbrain. (**B**) Immunostaining of DCX^+^/SOX2^+^ cells and pH3^+^ cells at day 28. Scale bars, 200 µm (upper panels) and 400 µm (lower panels). (**C**) Immunostaining of FOXA2^+^/TH^+^ cells and NURR1^+^/TH^+^ cells at day 28. Scale bars, 100 µm. (**D**) Violin plots of TH, LMO3 and ALDH1A1 generated from scRNA-seq data of developing human ventral midbrain. (**E** and **F**) qPCR analysis of LMO3 (**E**) and ALDH1A1 (**F**) at day 28. *p < 0.05 vs. vehicle (n = 3). (**G**) Gene expression analysis of differentiation protocol based on qPCR. Values are color coded and normalized to the sample with highest expression for each gene (n = 2-3).

### Analysis of hESC-derived cell types by scRNA-seq

To further examine the quality of the hESC-derived midbrain cell types, we performed single cell transcriptomics profiling of H9 and HS980 cells at days 0, 11, 16, 21 and 28 of differentiation. The quality of the cells was monitored by immunocytochemistry and found to be comparable for both cell lines at day 16 (**Figure S4A** and **S4B**).

After filtering, 11 681 high quality cells were included in our analysis. The mean UMIs of cells at different time-points of differentiation ranged between 6753 and 15482, and their mean transcripts between 2490 and 3931 (Figure 5A). Both H9 and HS980 contributed similar proportions of cells to the time-points and clusters **(Figure 5SA and S5B)**. Dimensionality reduction and Louvain clustering with Cytograph revealed 38 clusters (Figure 5A). Cluster 1 was formed by undifferentiated hESCs at days 0 and 11, while clusters 0 and 2-11 were mainly contributed by cells from day 11-16, which have a higher proliferation index and are enriched in progenitor markers such as *SOX2* (Figures 5B and S5C and **S5D**). Clusters 12-37 were mainly contributed by days 21 and 28 and contained all the cells enriched in the expression of neuronal markers such as *STMN2* and *MYTL1* (Figures 5B, 5C and **S5E**).

**Figure 5.**
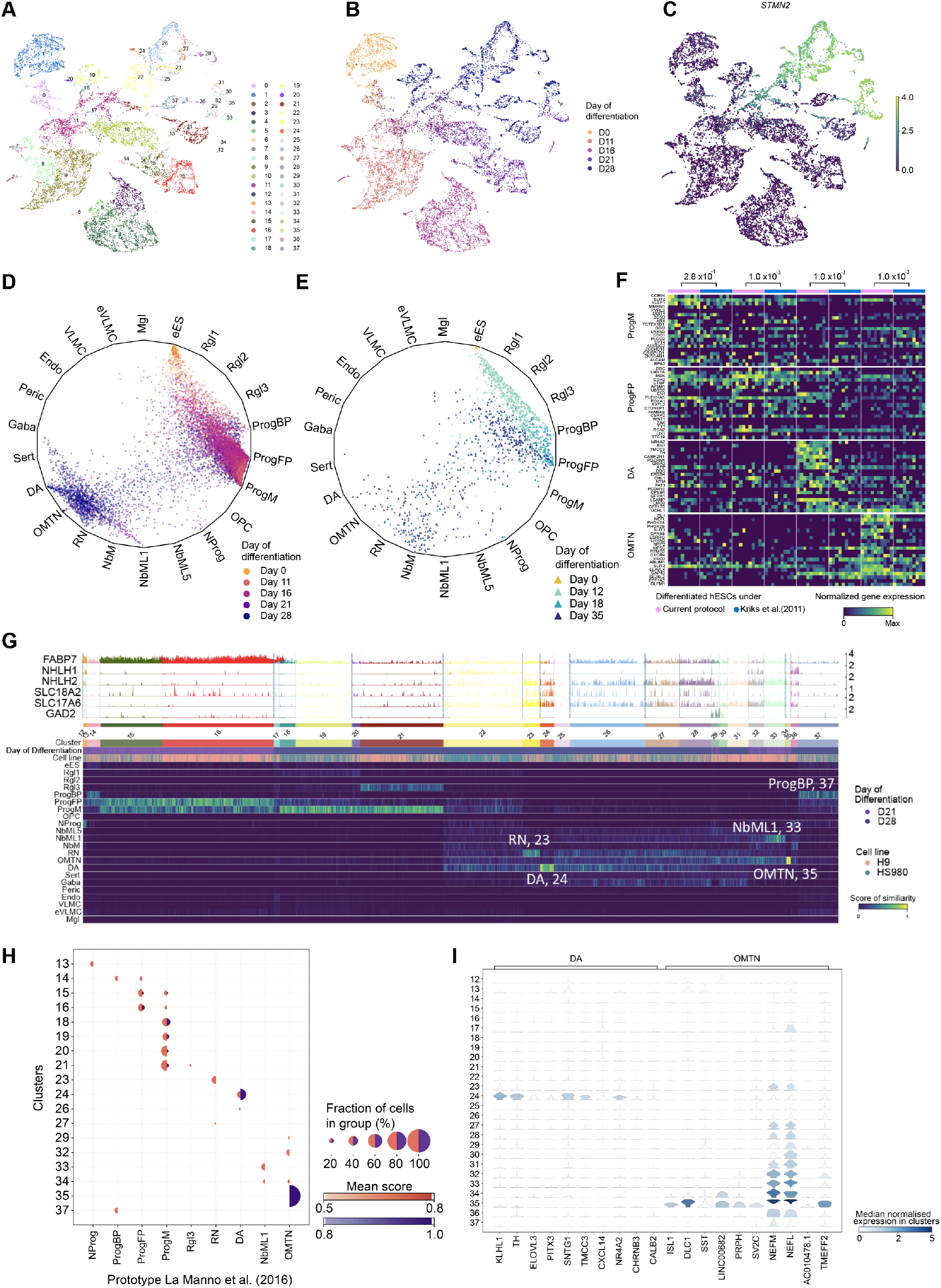
Analysis of hESC-derived cell types by scRNA-seq and logistic regression. (**A-C**) UMAP projection of hESC-derived cells after quality filtering showing cells coloured by their Louvain cluster’s membership (**A**), day of differentiation and analysis (**B**), and their log-library size normalised gene expression of *STMN2* (**C**). (**D** and **E**) Wheel plot showing a comparison by logistic regression of hESC-derived midbrain cell types (dots) generated by the protocol developed in this study (**D**) or the protocol by Kriks et al., (2011) (**E**), to endogenous human embryonic ventral midbrain cell types from La Manno et al., (2016) and vascular leptomeningeal cells from Marques et al., (2018). (**F**) Heatmap showing genes with highest coefficients from logistic regression for progenitor midline (ProgM), progenitor floor plate (ProgFP), dopaminergic neurons (DA) and oculomotor and trochlear nucleus (OMTNs). Log-library size normalized gene expression of top-similar hESCs to *in vivo* counterparts derived from current protocol and Kriks et al., (2011) are shown. Permutation test for each reference cell type with H_1_: The average of genes’ (shown on heatmap) expression sum of top-similar cells derived from current protocol was greater than top-similar cells derived from Kriks et al., (2011). Permutation was performed 1000 times on normalised counts, p-values are shown. (**G**) Track plot showing log-library size normalised gene expression of selected genes of cells at days 21 and 28 of differentiation (Top). Heatmap showing similarities between differentiated cells and reference endogenous cell types from La Manno et al., (2016) and Marques et al., (2018), scoring using logistic regression (Bottom). (**H**) Dot plot showing clusters from days 21 and 28 of differentiation with average similarity scores, as determined by logistic regression, between 0.5 and 0.79 (orange coloured) or between 0.8 and 1.0 (purple coloured). Only clusters have more than 0.5 similarity scores are shown. (**I**) Violin plot showing genes enriched in cluster 24 and 35. Log-library size normalized gene expression is shown.

### hESC derivatives are comparable to endogenous midbrain standards

Logistic regression was next used on scRNA-seq data to determine the probability of each of the hESC-derived cells being any of the endogenous human ventral midbrain tissue reference cell types as defined by La Manno et al. (2016) (Figure 6A-6C). In addition, we also used a reference dataset of vascular leptomeningeal cells (VLMCs) (Marques et al., 2018) because this cell type was not previously found in the endogenous developing human ventral midbrain in *vivo* (La Manno et al., 2016), but has been detected in hESC-derived midbrain cultures (Tiklová et al., 2019). We found that our human development-based hESC differentiation protocol generates cells with low or extremely low probability of being cell types defined by non-ventral midbrain standards, such as hindbrain serotonin neurons or VLMCs, respectively (Figure 5D). Consistent with this data, double COL1A1^+^ and PDGFRA^+^ VLMCs were not detected at day 16. However, early CHIR99021 treatment (days 0-2) and removal of high CHIR99021 (7.5 μM) in the presence of FGF8b (days 11-16) induced the emergence of strongly double-positive COL1A1^+^ and PDGFRA^+^ cells (**Figure 7**). These results show that VLMCs are not present in our standard midbrain culture conditions, but they can emerge by premature CHIR99021 treatment in the absence of CHIR99021 boost.

**Figure 6.**
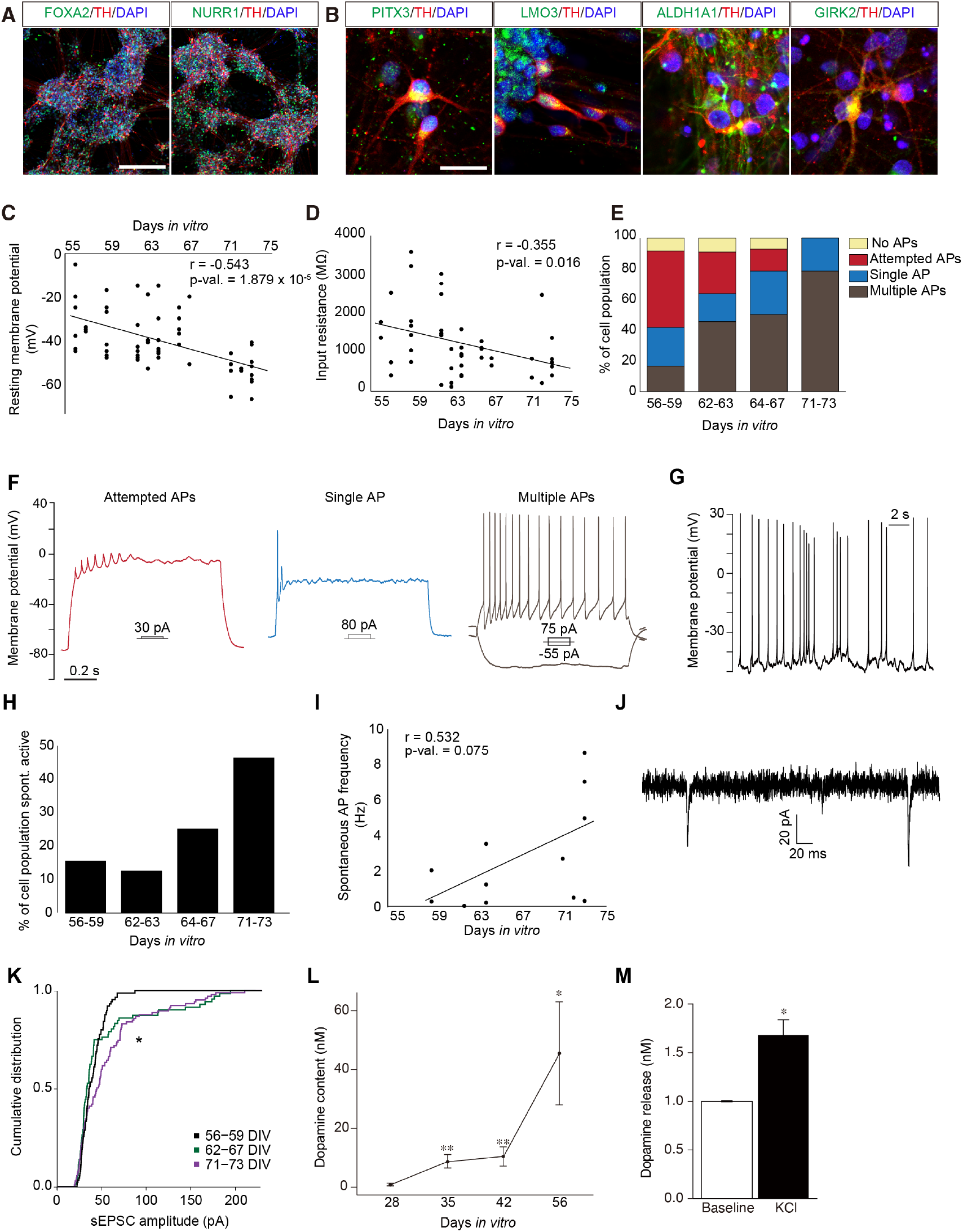
Maturation and functionality of the hESC-derived mDA neurons. (**A** and **B**) Immunocytochemical staining of FOXA2^+^/TH^+^ as well as NURR1^+^/TH^+^ (**A**), as well as PITX3^+^/TH^+^, LMO3^+^/TH^+^, ALDH1A1^+^/TH^+^, GIRK2^+^/TH^+^ (**B**) at day 56. Scale bar, 200 µm (**A**) and 25 µm (**B**). (**C-L**) Electrophysiological analysis of cells from days 56-73. (**C** and **D**) Decrease of resting membrane potential (**C**) (n = 55 cells) and reduction in input resistance (**D**) (n = 46 cells) with increasing days *in vitro* (DIV). (**E** and **F**) Percentage of cell population exhibiting the ability to generate the different spiking types in response to square current pulses as seen in example traces (**F**) (n = 12 cells for 56-59 DIV, n = 11 cells for 62-63 DIV, n = 14 cells for 64-67 DIV and n = 14 cells for 71-73 DIV). (**G**) Example trace of a spontaneous active neuron. (**H**) Percentage of each cell population spontaneously spiking (n = 13 cells for 56-59 DIV, n = 8 cells for 62-63 DIV, n = 12 cells for 64-67 DIV and n = 13 cells for 71-73 DIV). (**I**) Spontaneous action potential (AP) frequency of cells that were spontaneously spiking (n = 12 cells). (**J**) Example trace of a cell receiving two spontaneous excitatory post-synaptic currents (sEPSCs). (**K**) Cumulative distribution of sEPSC amplitudes in each cell population (Kolmogorov–Smirnov test: 56-59 DIV vs 62-67 DIV p-value = 0.243, 56-59 DIV vs 71-73 DIV p-value = 7.97 x 10^-4^, 62-67 DIV vs 71-73 DIV p-value = 9.67 x 10^-4^). (**L** and **M**) HPLC analysis of whole cell dopamine content from day 28-56 (**L**) and dopamine release at day 56 (**M**). *p < 0.05, **p < 0.01 (n = 4, error bars are S.E.M.).

We also found that cells generated in our cultures had high probability of being cell types defined by the *in vivo* human ventral midbrain standards (Figure 5D). The most abundant progenitor-like cell types were midline progenitors (ProgM) and floorplate progenitors (ProgFP), identified at day 16. Instead, postmitotic cell-types such as oculomotor and trochlear neurons (OMTNs) were found at day 21, and dopaminergic neurons (DA) at day 28. Notably, other neural cell types, such as oligodendrocyte progenitors or radial glia-like cells (Rgl2), both found in the basal plate (La Manno et al., 2016), were not found (Figure 5D). These findings indicate that the transcriptomic profiles of the hESC-derived cell types are comparable to that of cells in the most ventral aspect of the human midbrain *in vivo*.

### Improved quality and developmental dynamics of hESC-derived midbrain cell types compared to previous hESC differentiation conditions

Next we used our reference dataset to predict cell types from a previous scRNA-seq experiment (La Manno et al., 2016) in which H9 and HS401 hESC lines were differentiated for 12, 18 and 35 days into mDA neurons using the protocol by Kriks et al., (2011) (Figure 5E). Comparison of cells generated by the Kriks protocol to those generated by our new protocol with improved developmental control revealed four important differences. First, abundant basal plate progenitors but not midline progenitors were generated by the Kriks protocol (2011), while our new protocol generated more midline progenitors, reflecting an improved ventralization (Figure 5D and 5E). Second, a significant enrichment in the expression of genes defining endogenous human midline progenitors (ProgM), such as *CORIN, SLIT2, SULF1* and *ALCAM,* was found in the new compared to the old protocol (Figure 5F). Third, mDA neurons appeared at day 35 in the old protocol, but they were already abundant at day 28 in our new protocol (Figure 5D and 5E). Fourth, neurons of higher quality were found in the new protocol compared to the old one, as assessed by their similarity to the oculomotor (OMTN) and dopaminergic neuron (DA) standards (dots near the vertices in the wheel/polygon plot, Figure 5D and 5E). Moreover, genes typically expressed by OMTNs (i.e.: *ISL1, NEFL* and *PHOX2A*), or DA neurons (i.e.: *NR4A2, EN1* and *TH*) were significantly enriched in cells generated by the new protocol compared to the old one (Figure 5F). Combined, these results suggest that our new protocol, compared to that by Kriks et al. (2011), generates cultures with improved cell composition and cell types of a quality closer to that in the human ventral midbrain *in vivo*.

### Correct developmental dynamics and high quality cell types in hESC-derived cultures compared with endogenous midbrain standards

We first found that hESC-derived clusters enriched in day 21 and 28 cells (c12-37, Figures 5G and Fig**ure S6D**) are similar to several cell types in the endogenous human ventral midbrain from week 6-11 (La Manno et al., 2016). At day 21, clusters expressing high levels of *FABP7* (c12-16) contained cells that resembled either neuronal progenitors expressing the pro-neural genes *NHLH1* and *NHLH2* (NProg, c12-13) or both endogenous floorplate and midline progenitors (ProgFP and ProgM, c15-16), or both floorplate or basal plate progenitors (c14). Notably, hESC-derived postmitotic cells expressing *STMN2* and *MYT1L* were also found at day 21 (clusters 29, 32-35, Figure 5A-5C). These clusters resembled mediolateral neuroblasts (NbML1, c33) or oculomotor and trochlear neurons (OMTNs, c35) (Figure 5G), both of which appear early in the human ventral midbrain *in vivo* (La Manno et al., 2016).

At day 28, we also observed a cluster with progenitors similar to endogenous basal plate progenitors (ProgBP, cluster 37), but most of the progenitor cells resembled midline progenitors (ProgM, clusters 18-21), with cluster 21 exhibiting additional features of Rgl3 identity (Figure 5G). We found cells expressing neurogenesis markers such as *NHLH1* and *NHLH2* (c22), and markers such as SLC18A2 and/or SLC17A6 (clusters 26-28 and 30-31), expressed by nascent mDA neurons (Kouwenhoven et al., 2020). Moreover, postmitotic day 28 cells resembled endogenous human week 6-11 red nucleus neurons (RN, c23) and dopaminergic neurons (hDA, c24). Thus, our results indicate that the most prominent progenitors derived from hESCs are the floorplate and midline progenitors at day 21, followed by the midline progenitor at day 28. Notably, neurons are generated with a developmental timing similar to that of their *in vivo* counterparts, with OMTNs being detected at week 7 *in vivo* and day 21 *in vitro*, followed by DA neurons at week 8 *in vivo* and day 28 *in vitro*.

Next, we scored the degree of similarity between hESC-derived cells and the endogenous standards. Cells reaching the highest degree of similarity (score 0.8-1) in ascending order were: floorplate progenitors (ProgFP) (c15,16), midline progenitors (ProgM) (c18-21), mDA neurons (c24) and OMTNs (c35) (Figure 5H). Notably, 50% of the cells in c24 and 96% in c35 highly resembled endogenous DA neurons (mean similarity score of 0.91) and OMTNs (0.98), respectively (Figure 5H). Accordingly, c35 was found selectively enriched in genes that define OMTNs such as *ISL1* and *DLC1* (c35), while c24 was enriched in DA neuron genes, including *NR4A2*, *TH* and *PITX3* (Figure 5I). Combined, these results indicate that our current differentiation protocol sequentially generates good quality progenitors (floorplate followed by midline progenitors), and very high-quality neurons (OMTNs followed by mDA neurons), closely resembling endogenous midbrain development.

### hESC-derived mDA neurons become mature functional neurons

We first explored whether the dopaminergic neurons generated *in vitro* at day 28 can develop into mature dopaminergic neurons. Analysis of marker expression at day 56, revealed that most of the TH^+^ neurons are FOXA2^+^ and NURR1^+^ (Figure 6A). Moreover, some of the TH^+^ neurons were positive for markers associated to mature mDA neurons, such as PITX3^+^, LMO3^+^, ALDH1A1^+^ or GIRK2^+^ (Figure 6B), indicating that hESC-derived mDA neurons adopt mature midbrain phenotypes.

We next performed electrophysiological recordings to examine whether these cells can mature into functional neurons. Analysis of the membrane resting potential and input resistance revealed a progressive decrease of these two parameters from day 56 until day 73, indicating further maturation during this period (Figure 6C and 6D). Similarly, current-clamp recordings from day 56 to 73 revealed improved firing capacity upon current injections (Figure 6E and 6F). Notably, the proportion of neurons responding with multiple or single action potentials increased from 42% at days 56-59 to close to 100% at days 71-73. Moreover, the proportion of neurons exhibiting spontaneous electrical activity and action potentials also increased from day 56 to 73 (Figure 6G-6I). In addition, the patched cells received spontaneous excitatory postsynaptic currents (sEPSCs) from days 56 to 73 (Figure 6J), suggesting the establishment of synaptic connections. Combined, these results indicate that our human development-based differentiation protocol gives rise to neurons that progressively mature in culture and become electrophysiologically active by days 71-73. Finally, we examined whether these neurons also acquire the capacity to synthesize and release the neurotransmitter dopamine (Figure 6L and 6M). High performance liquid chromatography (HPLC) revealed very low levels of dopamine content at day 28, an increase at days 35 and 42 and high levels at day 56, whereas dopamine release was only detected at day 56. Thus, the results above show that hESC-derived mDA neurons progressively acquire functional properties of mature mDA neurons *in vitro*.

## Discussion

In this study we address the challenge of achieving hESC-derived products with sufficient molecular definition and similarity to endogenous standards to enable their future development for drug development and cell replacement therapy. We show that by modulating different pathways in a time-controlled manner in hESCs, it is possible to reproduce key aspects of the developmental dynamics of the ventral midbrain and improve midbrain patterning as well as mDA neurogenesis and differentiation. Indeed, scRNA-seq analysis revealed sequential generation of hESC-derived ventral midbrain progenitors and neurons with single cell transcriptomics profiles similar to those found in the endogenous human midbrain. Moreover these profiles were of higher quality than those obtained with a previous mDA differentiation protocol. In addition, we find that hESC-derived mDA neurons can mature and become functional *in vitro*. Indeed, mDA neurons appear by day 28, express mature mDA markers by days 42-56, acquire the capacity to release dopamine by day 56 and become electrophysiologically active neurons by day 73.

We hereby define the function of a number of key developmental pathways, which have not been previously examined or used to differentiate hESCs into mDA neurons. Factors such as full length LN511, the morphogen WNT5A, and the combination of FGF8b and high CHIR99021 (7.5 μM) were found to affect anterior-posterior patterning. For instance, LN511 and WNT5A decreased the expression of hindbrain genes such as *GBX2* and *HOXA2* at day 11. Moreover, WNT5A as well as high CHIR99021 combined with FGF8b, decreased the expression of anterior and lateral genes (*FOXG1, BARHL1, PITX2, SIX3* and *NKX2.1*), and increased expression of midbrain genes (*LMX1A* and *EN1*) at days 11 and 16. In addition, cell types such as VLMCs, absent in our developmental standards and in other hESC-derived ventral midbrain cultures (Kim et al., 2021), were not detected during mDA differentiation. Interestingly, a modified protocol involving early CHIR99021 administration (Day 0-11) and FGF8 treatment in the absence of CHIR99021 boost (Day 9-16) gave rise to VLMCs, suggesting this cell type emerges in specific culture conditions, as previously reported (Tiklová et al., 2020).

We also found that sequential administration of the small molecules CHIR99021 (7.5 μM) and GW3965, to activate Wnt/β-catenin and LXR signaling respectively, control different aspects of neurogenesis. Indeed, high CHIR99021 increased the number of NGN2^+^ cells at day 16, a gene required for mDA neurogenesis (Kele et al., 2006), while GW3965 improved neurogenesis (EdU^+^/DCX^+^ cells) and reduced the number of SOX2^+^ cells at day 21. Additionally, we found that treatment with the FGF receptor inhibitor, SU5402, and the MEK/ERK inhibitor, PD0325901, further reduced proliferation and SOX2^+^ cells at day 28. At this stage, abundant TH^+^ neurons expressed midbrain markers such as LMX1A, FOXA2, NURR1, PITX3, LMO3, ALDH1A1 and GIRK2 (KCNJ6), indicating efficient mDA neurogenesis.

Single cell transcriptomics allowed us to perform a detailed analysis of the molecular cell types generated in our hESC cultures compared with endogenous human ventral midbrain standards (La Manno et al., 2016). This comparison enabled us to define the identity of the cell types generated *in vitro* as well as their quality and their developmental dynamics. Analysis of hESC-derived cultures revealed the presence of good quality progenitors, which followed a temporal sequence of events similar to that found *in vivo*. The floorplate progenitor was enriched at day 21 and was nearly absent at day 28 (c14-16), whereas the midline progenitor (c18-21) was present at day 21 and is abundant at day 28. Notably we found that the identity of progenitors at day 21 was less well defined than at day 28. Indeed, day 21 progenitor clusters contained two types of progenitors, floorplate and basal plate (c14) or floorplate and midline (c15, 16). In addition some progenitors in these clusters partially shared the two identities, suggesting the presence of cell transitions or earlier progenitors that are not present in week 6-11 developmental standards and are thus only partially recognized. Instead, day 28 clusters contained only one type of progenitor, either midline (c18-21) or basal plate progenitors (c37), suggesting that they have refined their identities and are then recognized by our developmental standards.

As expected by the presence of basal plate, floorplate and midline progenitors, our cultures give rise to diverse postmitotic cell types found in the endogenous human ventral midbrain during week 6-11. Interestingly these cells also emerge following a specific developmental sequence of events, with mediolateral neuroblast 1 (NbML1, c33), and oculomotor and trochlear neurons (c35) emerging at day 21, followed by red nucleus (c23) and mDA neurons (c24) at day 28. Notably, the quality of mDA neurons was very high already at day 28, with 50% of the hESC-derived mDA neurons showing a transcriptome 91% similar to that of endogenous embryonic human mDA neurons. These results show that our human development-based differentiation protocol, by improving developmental control of hESCs during mDA differentiation, recapitulates multiple aspects of human ventral midbrain development, including the temporal axis and the generation of high-quality prototypical cell types defined by scRNA-seq analysis. We therefore suggest the current differentiation paradigm may be useful to model and study human mDA neuron development and functionality *in vitro*. Moreover, since human ventral midbrain tissue has been successfully used for cell replacement therapy in PD patients (Kefalopoulou et al., 2014; Li et al., 2016; Lindvall et al., 1990), and hPSC-derived DA progenitors are currently being used in clinical trials for PD cell replacement therapy (Barker et al., 2017; Doi et al., 2020; Kim et al., 2021; Piao et al., 2021; Schweitzer et al., 2020; Tao et al., 2021), we suggest our differentiation paradigm may also be useful for this type of application. We envision that strategies aiming at generating or selecting molecularly-defined cell-types, such as the progenitor of the dopaminergic neuron subtype mainly affected by disease, the SOX6_AGTR1 subpopulation (Kamath et al., 2022), may enable highly precise and safe cell replacement therapy for PD.

In the near future, we expect that hESC-derived preparations destined for cell replacement therapy will be routinely examined at the single cell level in order to control for cell composition and quality. In this context, our work represents a first attempt to compare cell preparations with endogenous standards, but more work will be needed to improve the resolution of this type of analysis. This will involve: i) improving the definition of endogenous human midbrain standards, with more time-points, deeper coverage and multimodal single cell data; ii) correlating cell composition and quality at the single cell level *in vitro* with the preclinical and clinical performance of the grafts; and iii) developing new computational methods and tools to integrate multiple levels of information and precisely compare hPSC-derived cell types with endogenous standards and functionality *in vitro* and *in vivo*. Ultimately, we should be able to design and develop hPSC preparations with the desired composition, single cell quality and functionality for specific and precise *in vitro* and *in vivo* applications.

### Experimental Procedures

#### Undifferentiated human ESC culture

Human ESC lines H9 (Thomson et al., 1998), HS401, HS975 and HS980 (Rodin et al., 2014) were maintained on LN521 (BioLamina)-coated dish in NutriStem XF hESC medium (Biological Industries). The cells were passaged with TrypLE Select (Thermo Fisher Scientific) every 4-6 days, and were replated at a density of 50,000-100,000 cells/cm^2^ in medium supplemented with 10 µM Y27632 (Tocris) for the first 24 hr.

### Differentiation protocol into mDA neurons

The cells were seeded at a density of 500,000 cells/cm^2^ on LN511 (BioLamina)-coated dish in NutriStem XF hESC medium with 10 µM Y27632. The cells were differentiated in TeSR-E6 medium (Stem Cell Technologies) supplemented 0.1 mM nonessential amino acids (Thermo Fisher Scientific), L-glutamine (Thermo Fisher Scientific), and 0.1 mM 2-mercaptoethanol (Gibco). 200 nM LDN193189 (Stemgent), 10 µM SB431542 (Tocris) and 10 µM Y27632 were supplemented from day 0, 2 µM purmorphamine (Stemgent) was supplemented from day 1, and 1.5 µM CHIR99021 (Sigma) was supplemented from day 3. 10 µM Y27632 was removed from culture medium on day 3. The medium was gradually changed to neurobasal medium (Thermo Fisher Scientific) with B27 supplement (Thermo Fisher Scientific) and 2 mM L-glutamine from day 5 to day 11. 100 ng/mL Wnt5A (R&D Systems) was supplemented from day 7. 100 ng/mL FGF8b (Peprotech) was supplemented from day 9. On day 11, the cells were dissociated into single cells and were replated on LN511-coated dish at a density of 500,000 cells/cm^2^ in neurobasal medium with B27 supplement and 2 mM L-glutamine. 100 ng/mL FGF8b, 7.5 µM CHIR99021 were supplemented from day 11 to day 16. 10 µM Y27632 was supplemented in medium first 24 hr after replating. On day 16, the cells were dissociated into single cells and were replated on LN511-coated dish at a density of 700,000 cells/cm^2^ in neurobasal medium with B27 supplement and 2 mM L-glutamine. 10 µM GW3965 (Sigma), 10 µM DAPT (Sigma), 20 ng/mL brain-derived neurotrophic factor (BDNF) (R&D Systems), 200 µM ascorbic acid (Sigma) were supplemented from day 16 to day 21. 10 µM Y27632 was supplemented in medium first 24 hr after replating. 10 µM GW3965 was removed from culture medium on day 21. 1 µM PD0325901 (Sigma), 5 µM SU5402 (Sigma), 10 ng/mL glial-cell derived neurotrophic factor (GDNF) (R&D Systems), 500 µM dbcAMP (Sigma) and 1 ng/mL transforming growth factor (TGF)β3 (R&D Systems) were supplemented from day 21 and PD0325901 andSU5402 were removed from day 28 (Figure 1A).

### Derivation of VLMCs from hESCs

Different variables in our mDA differentiation protocol were modified to examine whether VLMCs can be generated from hESCs when differentiated in culture conditions found in other protocols in the literature (Doi et al., 2014; Kim et al., 2021; Tiklová et al., 2019). To this end we compared administration of CHIR99021 (1.0 µM) at day 0 vs day 2 (1.5 µM, our protocol), which we suspected may play a key role, and combined these two variables together with FGF8b (100 ng/mL in our protocol vs no treatment), CHIR99021 at day 11 to day 15 (7.5 µM in our protocol vs no treatment), as well as administration of growth factors and small molecules from day 11 to 15 (20 ng/mL BDNF, 20 ng/mL GDNF, 500 µM dbcAMP and 200 µM AA). Cells were replated at day 16 in neurobasal medium with B27 supplement, 2 mM L-glutamine, 100 ng/mL FGF8b, 20 ng/mL BDNF, 20 ng/mL GDNF, 10 µM DAPT, 500 µM dbcAMP, 200 µM AA and 10 µM Y27632. Analysis was performed by immunocytofluorescence at day 17.

### cDNA synthesis and qPCR

Total RNA was extracted from the cells using RNeasy Plus Kit (Qiagen). 500 ng-1 µg total RNA was used for reverse transcription by a Super Script II First strand synthesis system with random primer (Thermo Fisher Scientific). qPCR was performed by using StepOne detection system (Applied Biosystems). Data analysis is based on dCt method with normalization of the raw data to *GAPDH* genes. Primer sequences were listed in Supplementary Table 1.

### Fluorescent immunocytochemistry

The cells were fixed by 4% paraformaldehyde (PFA) for 30 min at 4°C. The samples were then pre-incubated by 5% donkey serum in phosphate buffered saline containing 0.3% Triton X-100 (PBST) for 1 hr. The samples were incubated with primary antibodies at 4°C overnight. The samples were incubated with either Alexa488 or Alexa555 or Alexa647-conjugated secondary antibodies (Thermo Fisher Scientific) for 30 min and then were incubated with 4’,6-diamidino-2-phenylindole (DAPI) for 15 min. The primary antibodies were used as follows: ALDH1A1 (rabbit, 1:1,000, Abcam, ab23375), COL1A1 (sheep, 1:200, R&D Systems, AF6220), CORIN (rat, 1:1,000, R&D Systems, MAB2209), DCX (goat, 1:500, SantaCruz, sc-8066), EN1 (mouse 1:50, DSHB, 4G11), FOXA2 (goat, 1:500, R&D Systems, AF2400), GIRK2 (rabbit, 1:400, Alomone, APC006), LMX1A (rabbit, 1:4,000, Millipore, AB10533), LMO3 (goat, 1:200, SantaCruz, sc-82647), MAP2 (mouse 1:1,000, Sigma, M4403), NGN2 (goat, 1:200, SantaCruz, sc-19233), NURR1 (rabbit, 1:500, SantaCruz, sc-990), OTX2 (goat, 1:1,000, R&D Systems, AF1979), pH3 (rabbit, 1:500, Millipore, 06-570), PITX3 (goat, 1:500, SantaCruz, sc-19307), PDGFRa (rabbit, 1:100, Cell Signaling, 5241), SOX2 (rabbit, 1:500, Millipore, AB5603), TH (rabbit, 1:1,000, Millipore, AB152), TH (mouse, 1:500, ImmunoStar, 22941), and TH (sheep, 1:500, Novus, NB300).

### EdU pulse and chase

Click-iT EdU Imaging Kit (Thermo Fisher Scientific) was used for EdU pulse and chase experiment.10 µM EdU was supplemented into the culture medium for 4 hr at day 16 of differentiation, and then the cells were cultured until day 21. After fixation by 4% PFA, EdU detected was performed according to the manufacturer’s protocol.

### Single-cell RNA-sequencing libraries preparation

hESCs and cells of differentiation days 11, 16, 21, 28 were thawed and resuspended in NeutriStem medium with 10 μM Y27632. Cells were centrifuged at 300 x g for 2 min. Cell pellets were resuspended in BD staining buffer (FBS) at room temperature. Cell suspension was mixed with Sample Tag and incubated at room temperature for 20 min. Cells were washed with BD staining buffer twice and resuspended in PBS (with 0.04% BSA). Cells were filtered and counted. For each hESC line, we pooled 1000 cells from day 0, 1200 cells from differentiation days 11, 16 and 21, and 1400 cells from day 28. The pooled cells were processed with single cell capture, reverse transcription, and cDNA amplification according to the 10x Chromium^TM^ Single Cell 3’ Reagent Kits v2 User Guide. The corresponding Sample Tag libraries were prepared according to BD Single-Cell Multiplexing Kit—Human.

### Processing of single cell data

The samples were aligned to a combined reference genome of GRCh38.p12 and BD sample tags using Cellranger v3.0.2. Samples were then demultiplexed using BD Genomics Sample Multiplexing tools v0.4 from Cellranger outputs. Final UMI count matrices were obtained by running Velocyto v0.17 with default parameters for Chromium 10X samples on demultiplexed bam files with a gtf file combined of the human reference and BD sample tags. Data was filtered and processed using Cytograph^32^ with the following quality parameters. For each cell line count matrices, cells with less than 2000 UMIs detected transcripts, more than 5% of mitochondrial genes in their library, and identified as doublets by the Cytograph version of DoubletFinder, were excluded from further analysis. After filtering, there were 1359, 2372, 2714, 1962, 3274 cells remained at day 0, day 11, day 16, day 21 and day 28 of differentiation respectively with mean UMIs between 6 753 and 15 482, and mean detected transcripts between 2 490 and 3 931 (Figure 5A). Dimension reduction and Louvain clustering were then performed using Cytograph with PCA using 40 components on highly variable genes detected by variances. Clusters were refined by aggregating clusters that were not transcriptionally differentiated by manual inspection of gene enrichment for each cluster, resulting in 29 clusters (Figure 5B). Then, cells from days 21 and 28 of differentiation (cluster 11 onwards) were pulled for another iteration of processing as described. Final cluster membership (Figure 5A) was obtained by aggregating clustering results from the iterations.

### Comparison of hESC-derived cells to the human ventral midbrain development reference dataset

UMI matrices of hESCs-derived cells from the current protocol were log-transformed, normalized and scaled to the reference data with selected gene set (described below). The same transformation was performed to UMI matrices from previous hESC-derived midbrain cells (La Manno et al., 2016), which were differentiated as described (Kriks et al., 2011). The similarities of *in vitro* cells to *in vivo* reference were measured as probabilities being each reference cell types, using logistic regression (described below) and visualized in a wheel plot as described (La Manno et al., 2016). In brief, the similarities to reference cell types are summarized as dot products to respective coordinates of reference cell types in the wheel, such that the distance to each reference cell type of an individual cell is in proportion to its relative similarity to the reference. For comparing the hESC-derived cells to the *in vivo* reference, L2-regularised logistic regression was used on human ventral midbrain cell types (La Manno et al., 2016), as well as mouse pericyte lineage cells (PLCs) and vascular leptomeningeal cells (VLMCs)(Marques et al., 2018). Logistic regression was implemented as described (La Manno et al., 2016), using the following prototypes: Embryonic stem cells (eES) consisting of eSCa, eSCb and eSCc; Floor plate progenitors (ProgFP) consisting of ProgFPM and ProgFPL; Radial glia 2 (Rgl2) consisting of hRgl2a, hRgl2b, hRgl2c; Dopaminergic neurons (DA) consisting of hDA0, hDA1 and hDA2; GABAergic lineage (Gaba) consisting of hGaba and hNbGaba; VLMCs consisting of pnVLMCs and VLMCs; Pericytes (Peric) consisted of hPeric and PLCs.

### Logistic regression training

For training the logistic regression, UMI matrices corresponding to cells belonging to cell types of interests (La Manno et al., 2016; Marques et al., 2018) were aggregated, after conversion of mouse genes to their human counterparts with Ensembl BioMart tool (GRCh37 version). Then a gene set (excluding sex, mitochondrial, erythrocytes and cell-cycle related genes) was selected from recursive feature elimination (RFE)(Guyon et al., 2002) using linear supper vector classification (SVC) with 5-fold stratified cross validation, step = 0.1, and F1-weighted score as the evaluation on log-transformed, max-normalized and median total UMIs-scaled data. The implementation was performed using the python package scikit-learn v0.23.1. With RFE, we have converged to a compact set of genes (n = 1184) that was discriminant between reference cell types, by eliminating incrementally 10% of the genes that were least important. To attest the performance of the selected gene set on cell type prediction, 80% of the normalized and scaled data was used for optimizing the regularization strength (C) with either the selected gene set or size-matched random gene set (n = 1184). For each condition, C was scanned using Optuna (Akiba et al., 2019), a Bayesian hyperparameter optimization, with the default sampler over log uniform distribution (between the range of 0.001 and 2) for 100 trials. Over the trials, C was optimized by evaluating the mean brier score of 5-folds train-test split per trial. Mean accuracy per trial was also logged (Figure 6A). The selected gene set was validated by using the optimized C to predict the 20% of the data at the end of the optimization (**Figure S6B** and **6C**). Final model with the full dataset was trained in the same approach (C = 1.99) and used for comparing *in vitro* cells.

### Analysis of dopamine release and content by HPLC

For dopamine measurement experiments, hESC-derived mDA progenitors were plated on LN511 coated 12-well plates at a density of 5 x 10^5^ cells/cm^2^ on day 16 and collected on days 28, 35, 42 and 56 of differentiation for dopamine contents and release. Cells were incubated in 200 μL of Neurobasal + N2 medium for 30 min at 37°C and the supernatant was collected. Cells were then incubated in Neurobasal + N2 medium supplemented with 56 mM KCl for 30 min at 37°C and the supernatant collected. The supernatant was immediately stabilized with 20 μL of 1 M perchloric acid containing 0.05% sodium metabisulphite and 0.01% ethylene-diamine-tetra-acetic acid (EDTA) disodium salt and the samples stored at −80°C. Two days later, cells were collected to measure intracellular dopamine content. On the day of analysis, samples were centrifuged at 16,000 x g for 10 min at 4°C and then filtered through 0.2 µm nylon membrane inserts by centrifugation at 4,000 x g for 5 min at 4°C. The HPLC-ECD system used was a Dionex Ultimate 3000 series (Dionex, ThermoFisher Scientific, USA) and the injection volume was 20 µL for each sample. Analyte separation was performed on a Dionex C18 reversed-phase MD-150 3.2 mm x 250 mm column (3 µm particle size). Column and analytical cell were kept at 30°C and the first and second analytical cell were set to −100 mV and +300 mV, respectively. The mobile phase was pumped at a flow rate of 0.4 mL/min and consisted of 75 mM monobasic sodium phosphate, 2.2 mM 1-octanesulfonic acid sodium salt, 100 µL/L triethylamine, 25 µM EDTA disodium salt and 10% acetonitrile (v/v), pH 3.0 adjusted with 85% phosphoric acid. Chromatograms were acquired with Chromeleon software (Dionex, ThermoFisher Scientific) over an acquisition time of 55 min. Dopamine concentration was calculated for each sample. For dopamine release experiments, dopamine levels after KCl stimulation were normalized to unstimulated dopamine levels. For intracellular dopamine content, dopamine content was normalized to intracellular dopamine content on day 28. Data is shown as averaged normalized values from three independent experiments.

### Electrophysiological recordings

For patch-clamp electrophysiological recordings, hESC-derived mDA progenitors were plated on LN511 coated 24-well plates at a density of 500 000 cells/cm^2^ on day 16 and recordings were performed between days 56-73 of the differentiation protocol. Cells with neurites and non-flat cell body, general morphological aspects of neurons, were selected for whole-cell patch clamp recordings. Borosilicate glass pipettes (4-10 MΩ) were filled with intracellular solution containing 105 mM K-gluconate, 30 mM KCl, 10 mM Na-phosphocreatine, 10 mM HEPES, 4 mM Mg-ATP, 0.3 mM Na-GTP, and 0.3 mg/mL of Lucifer yellow (Sigma-Aldrich) (pH 7.3 adjusted with KOH). The cells were continuously perfused with a solution containing 140 mM NaCl, 2.5 mM KCl, 1.2 mM NaH_2_PO_4_, 1 mM MgSO_4_, 1.3 mM CaCl_2_, 10 mM glucose and 10 mM HEPES, pH 7.4, bath kept 30-35°C. The signal was amplified and digitized with Multiclamp 700B (Molecular Devices) and Digidata 1550 (Molecular Devices), respectively. Clampfit 11.1 (Molecular Devices) was used for analysis. For sEPSC detection, cells were clamped to −70 mV in voltage-clamp mode and 10-30 seconds were analyzed per cell by using template search (template made by averaging 10 typical EPSCs), false positives were manually removed by an analyzer blinded for the DIV of the cells. The remaining recordings were in current-clamp mode. Input resistances were calculated with a 1 second long ± 4-20 pA current injection. Spontaneous AP frequency was assessed during 30-60 seconds gap free recording. For spiking properties, cells that had a resting membrane potential above spiking threshold were brought down to ca −70 mV before current step injections. Pearson’s correlation (r) and two-tail p-values were reported.

### Statistic analysis

Results are given as means ± standard deviation (SD) or standard error of the mean (SEM). The significance of differences was determined by Student’s *t*-test for single comparisons and by one-way analysis of variance (ANOVA) or two-way ANOVA for multiple comparisons. Further statistical analysis for *post hoc* comparisons was performed using Tukey’s test (Prism 6; GraphPad, San Diego, CA, USA). Mann-Whitney U-test was also performed using R v4.1.1 to analyze the numbers of VLMCs. Permutation test was performed using python v3.7.7 for analyzing the average gene expression for top-similar cells as shown from logistic regression analysis between current protocol and the protocol by Kriks et al., 2011.

## Supporting information

Supplemental Figures 1-7 and Tabe 1

## Acknowledgments

We thank members of the Arenas lab for help, suggestions, and helpful discussions; Natalie Welsh for feedback on the manuscript; and BioLamina for providing laminins. This work was supported by Vetenskapsrådet (VR 2016-01526 and 2020-01426), European Commission grants Neurostemcell-repair (FP7, 602278) and Neurostemcell-reconstruct (H2020, 874758), ERC advanced grant (884608), Knut and Alice Wallenberg Foundation (KAW scholar 2018.0232), Karolinska Institutet StratRegen (SFO 2018), Cancerfonden (CAN 2016/572), Parkinsonfonden (900/16) and Hjärnfonden (FO2019-0068) to EA; by the Chan Zuckeberg Initiative and the Silicon Valley (2018-191929) to EA, SL and PS; by the Swedish Foundation for Strategic Research (SSF, SB16-0065) to EA and SL; by the Knut and Alice Wallenberg Foundation (2018.0172, 2018.0220), Erling-Persson Foundation (HDCA), and European Union (BRAINTIME) to SL. KN was supported by the Uehara Memorial Foundation, the Kyoto University Foundation, the Mochida Memorial Foundation for Medical and Pharmaceutical Research (6-2) and the Scandinavia-Japan Sasakawa Foundation (15-18); SY by Hjärnfonden (PS2018-0043); ESA by a KID grant (2-5996/2018).

## Author contributions

K. Nishimura, SY and EA designed the project. K. Nishimura performed the experiments in Figures 1-4, 6A, B and S1-S3. SY and ESA performed additional DA differentiation experiments (Figures 5 and 6) with support from CS and GL. SY and LH performed scRNA-seq with support from SL. KL performed the bioinformatic analysis in Figures 5, S5 and S6. K. Nikouei, SG and JHL contributed to the electrophysiological analysis and interpretation in Figure 6C-K. WP and PS contributed to the analysis of dopamine content and release in Figures 6L-M. EA supervised the project, co-wrote the manuscript with K. Nishimura, SY and KL. All authors reviewed and approved the manuscript.

