## Supplemental Figures 1-7 and Tabe 1 for "Single cell transcriptomics reveals correct developmental dynamics and high-quality midbrain cell types by improved hESC differentiation"

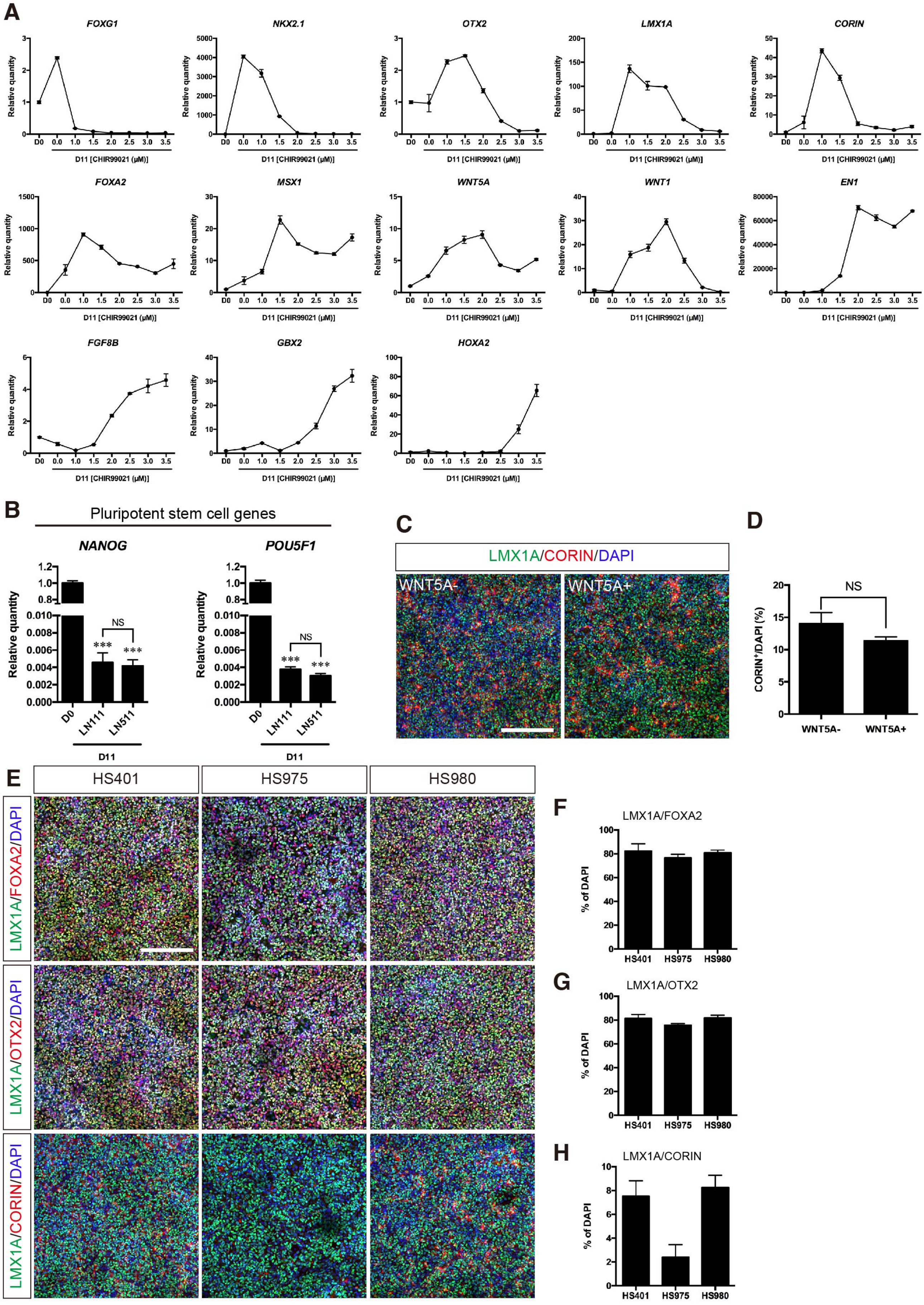

A
FOXG1
NKX2.1
OTX2
LMX1A
CORIN
FOXA2
MSX1
WNT5A
WNT1
EN1
FGF8B
GBX2
HOXA2
Relative quantity
D0
D11 [CHIR99021 (μM)]
B
Pluripotent stem cell genes
NANOG
POU5F1
Relative quantity
NS
***
LN111
LN511
D11
C
LMX1A/CORIN/DAPI
WNT5A-
WNT5A+
D
NS
CORIN+/DAPI (%)
WNT5A-
WNT5A+
E
HS401
HS975
HS980
LMX1A/FOXA2/DAPI
LMX1A/OTX2/DAPI
LMX1A/CORIN/DAPI
F
LMX1A/FOXA2
% of DAPI
G
LMX1A/OTX2
% of DAPI
H
LMX1A/CORIN
% of DAPI

**Figure S1. Patterning to floor plate progenitors from hESCs, related to Figure 1** (**A**) qPCR analysis of differentiated cells according to CHIR99021 concentration at day 11 (n = 3). (**B**) qPCR analysis of *NANOG* and *POU5F1* of differentiating cells on LN111 and LN511. ***p < 0.001 vs. D0. NS, not significant (n = 6). (**C**) Immunostaining of LMX1A$^+$/CORIN$^+$ cells at day 11. Scale bar, 200 μm. (**D**) Quantification of CORIN$^+$ cells at day 11. NS, not significant (n = 3). (**E**) Immunostaining of LMX1A$^+$/FOXA2$^+$ cells, LMX1A$^+$/OTX2$^+$ cells and LMX1A$^+$/CORIN$^+$ cells in HS401, HS975 and HS980 hESC lines at day 11. Scale bar, 200 μm. (**F**-**H**) Quantification of LMX1A$^+$/FOXA2$^+$ cells (**F**), LMX1A$^+$/OTX2$^+$ cells (**G**) and LMX1A$^+$/CORIN$^+$ cells (**H**) at day 11 in the WNT5A+ condition (n = 3).

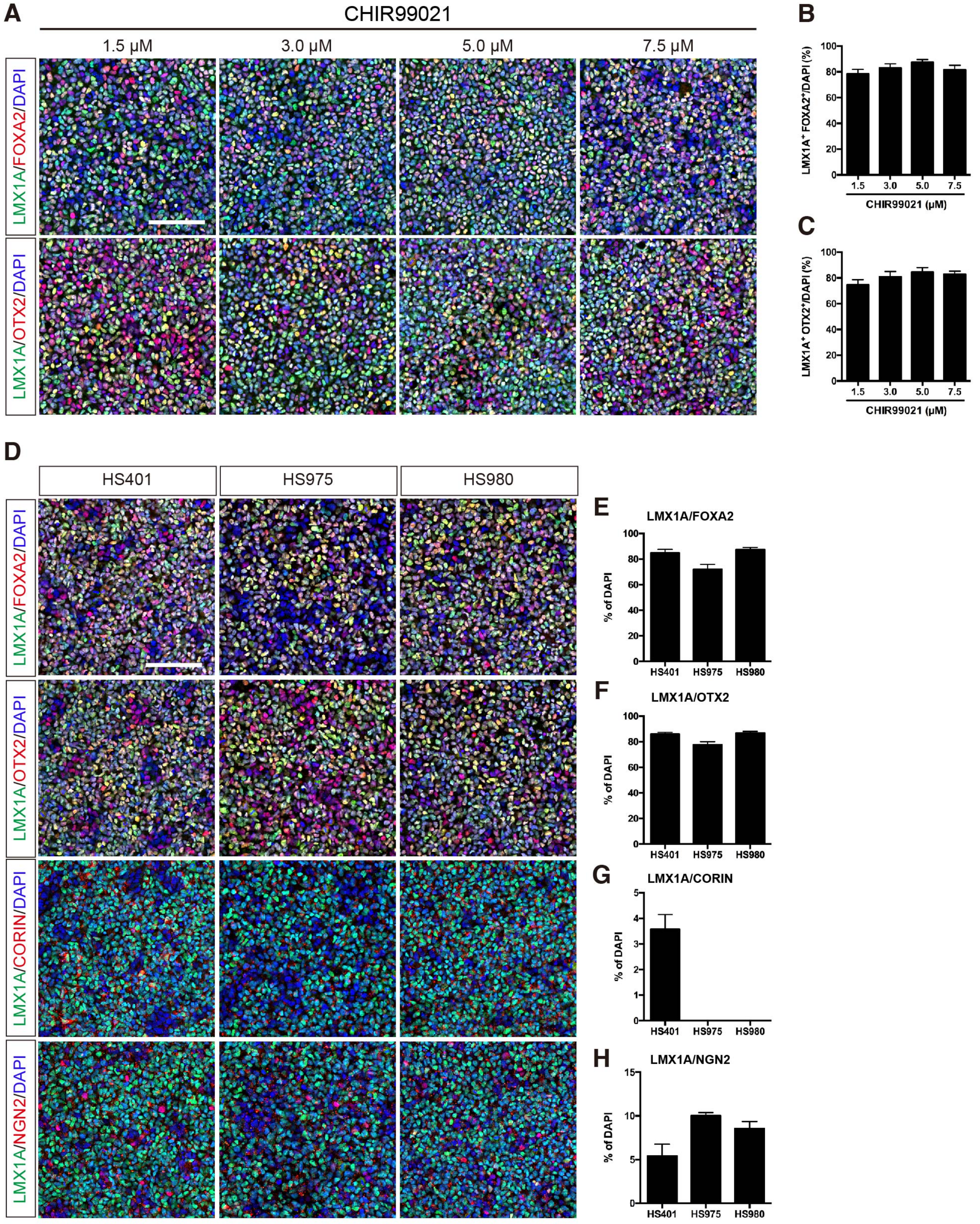
A
CHIR99021
1.5 μM
3.0 μM
5.0 μM
7.5 μM
LMX1A/FOXA2/DAPI
LMX1A/OTX2/DAPI
B
LMX1A+ FOXA2+/DAPI (%)
CHIR99021 (μM)
C
LMX1A+ OTX2+/DAPI (%)
CHIR99021 (μM)
D
HS401
HS975
HS980
LMX1A/FOXA2/DAPI
LMX1A/OTX2/DAPI
LMX1A/CORIN/DAPI
LMX1A/NGN2/DAPI
E
LMX1A/FOXA2
% of DAPI
F
LMX1A/OTX2
G
LMX1A/CORIN
H
LMX1A/NGN2

**Figure S2. Characterization of DA progenitors derived from three different hESCs, related to Figure 2.** (**A**) Immunostaining of LMX1A$^+$/FOXA2$^+$ cells and LMX1A$^+$/OTX2$^+$ cells at day 16, after exposure of HS980 cells to different concentrations of CHIR99021. Scale bar, 100 μm. (**B** and **C**) Quantification of LMX1A$^+$/FOXA2$^+$ cells (**B**), LMX1A$^+$/OTX2$^+$ cells (**C**) at day 16 (n = 3). (**D**) Immunostaining of LMX1A$^+$/FOXA2$^+$ cells, LMX1A$^+$/OTX2$^+$ cells, LMX1A$^+$/CORIN$^+$ cells and LMX1A$^+$/NGN2$^+$ cells in HS401, HS975 and HS980 hESC lines at day 16 of the development-based differentiation protocol with 7.5 μM CHIR99021. Scale bar, 100 μm. (**E**-**H**) Quantification of LMX1A$^+$/FOXA2$^+$ cells (**E**), LMX1A$^+$/OTX2$^+$ cells (**F**), LMX1A$^+$/CORIN$^+$ cells (**G**) and LMX1A$^+$/NGN2$^+$ cells (**H**) at day 16 (n = 3).

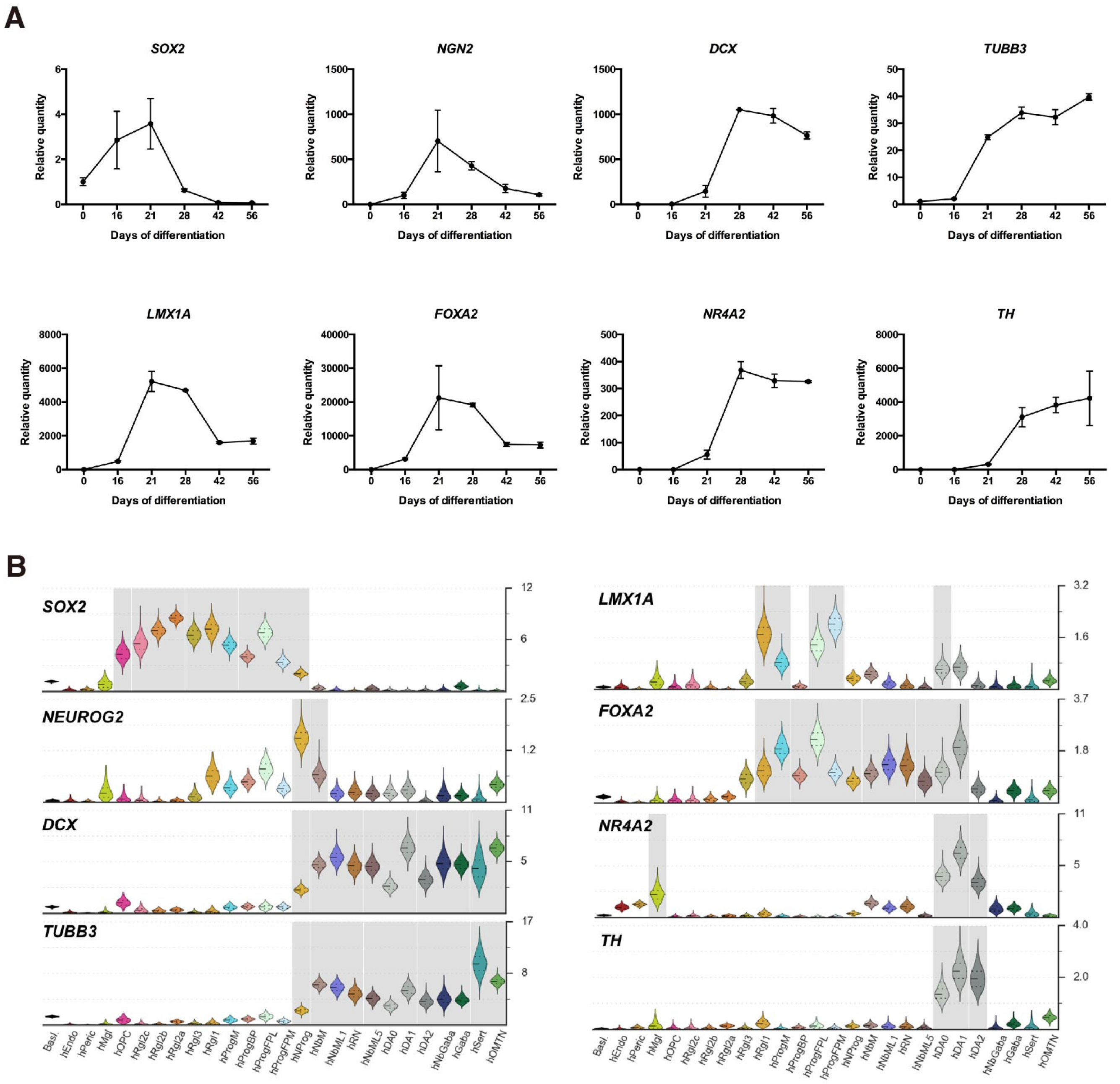


**Figure S3. Time course of key midbrain gene expression during hESC differentiation into mDA neurons, related to Figure 4.** (**A**) qPCR analysis of neural and neuronal markers and mDA neuron markers (n = 3) (**B**). Violin plots of general neural/neuronal markers and midbrain DA neuron markers generated from scRNA-seq data of developing human ventral midbrain.

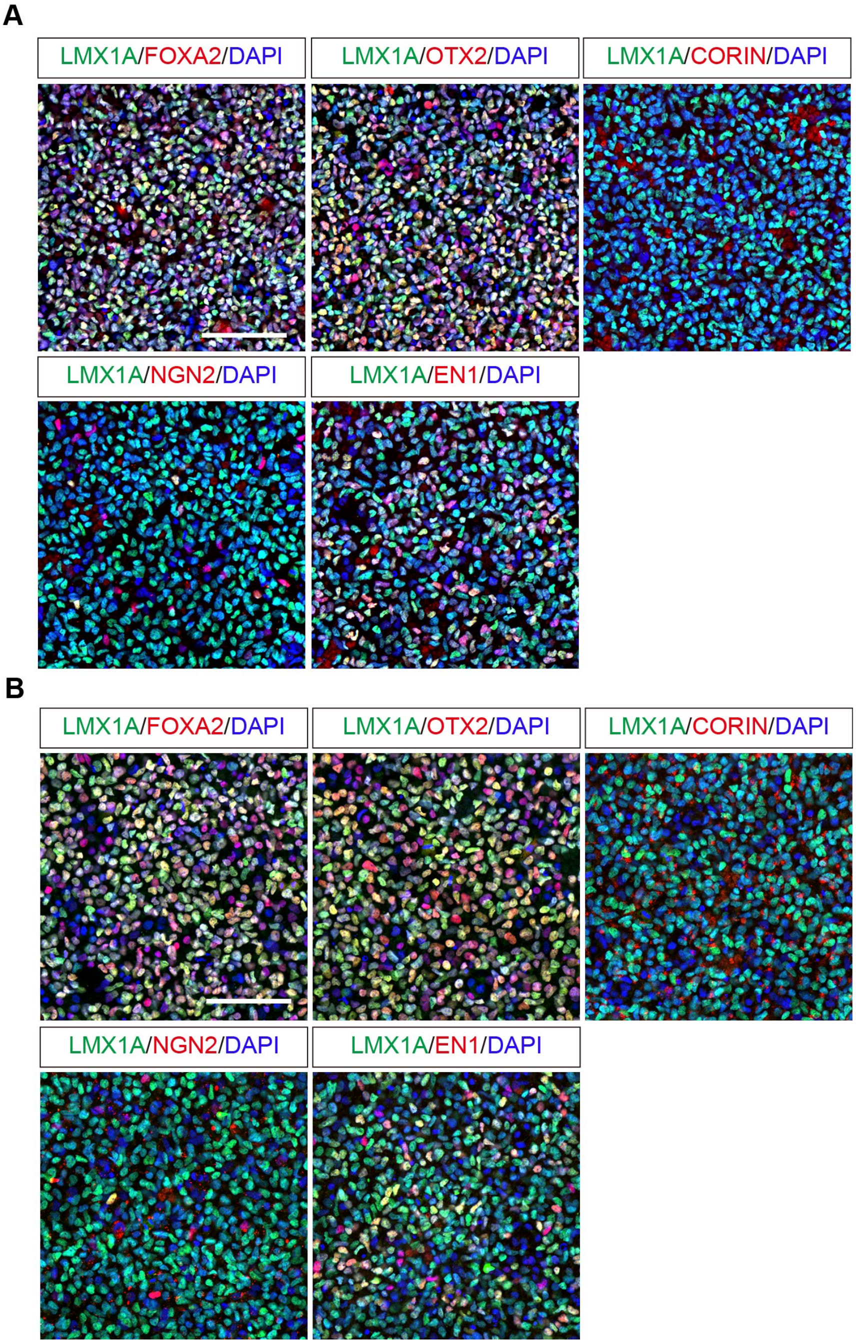


**Figure S4. Validation of hESC-derived mDA progenitors by immunocytochemistry prior to scRNA-seq, related to Figure 5.** Immunofluorescence of hESC-derived mDA progenitor-derived from H9 (**A**) and HS980 (**B**) hESCs at day 16. Scale bar, 100 μm.

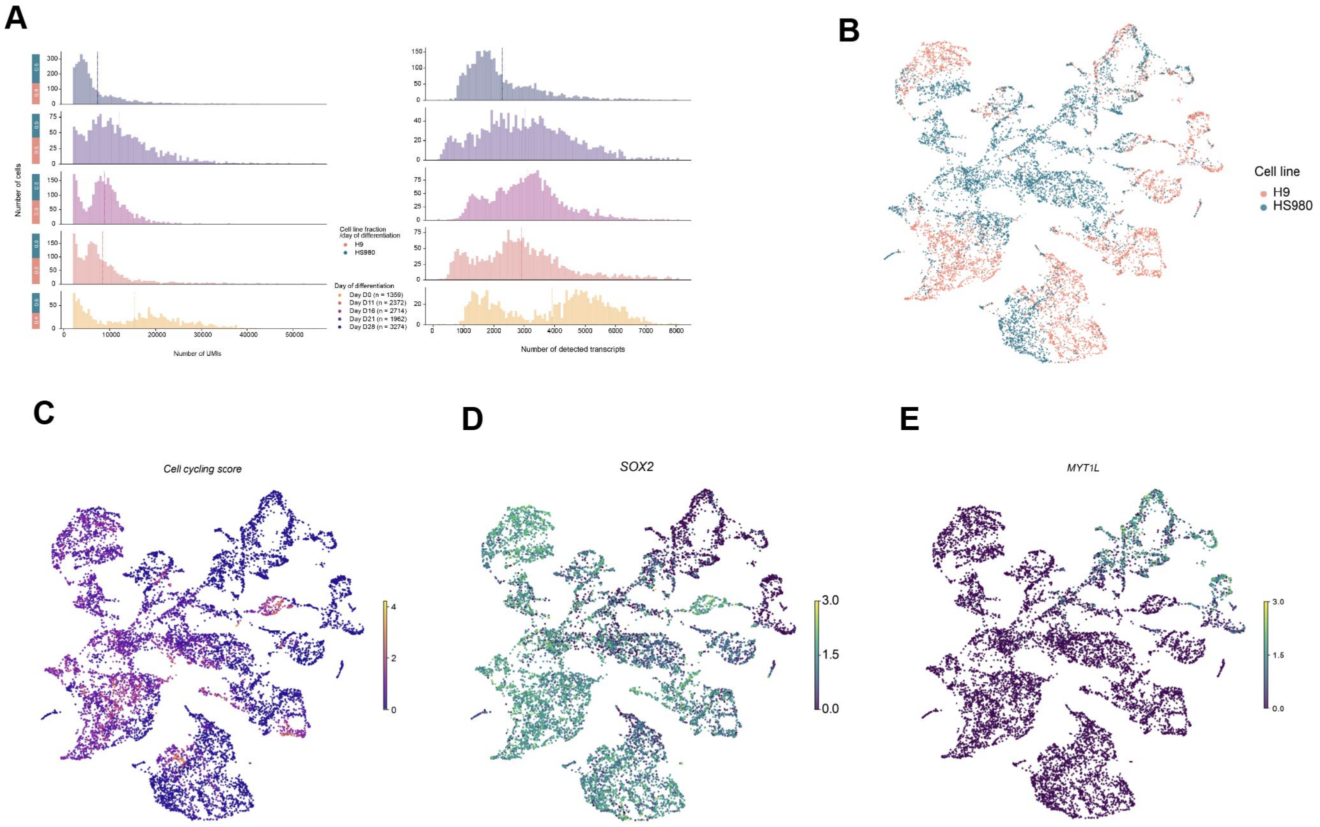


**Figure S5. Characterization of the sequenced hESCs-derived cells, related to Figure 5.** (**A**) Histogram showing the distribution of UMIs (right panel) and detected transcripts (left panel) per cell and day of differentiation. Bars show the fractions of cells generated by each of the two cell lines (H9 or HS980) at the indicated day of differentiation. (**B-E**) UMAP projection of hESCs-derived cells as in Figure 5A, showing cells coloured by cell line of origin (**B**) cell cycle score (**C**) and their log-library size normalised gene expression of *SOX2* and *MYT1L*.

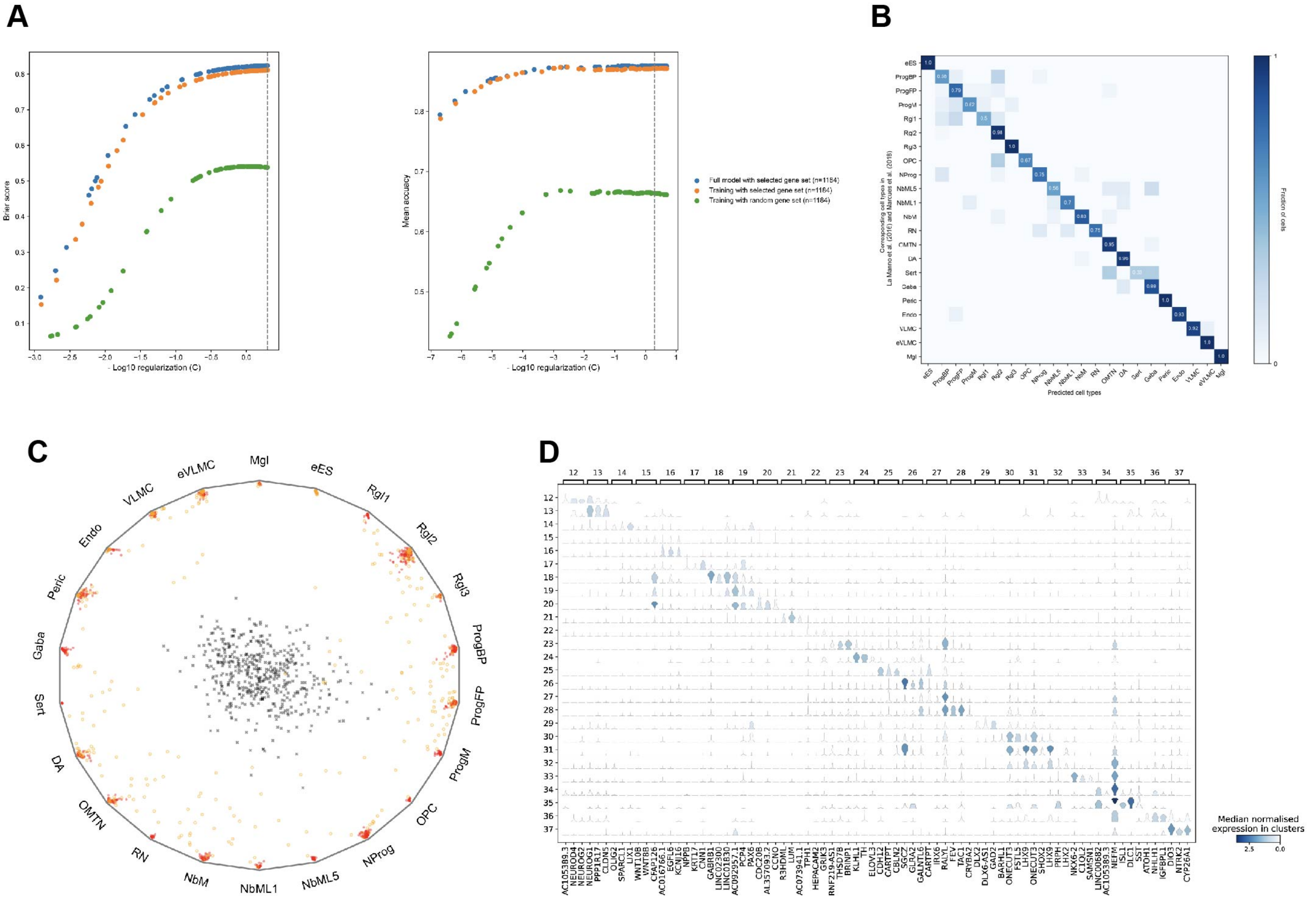


**Figure S6. Logistic regression training and cluster analysis, related to Figure 5.** (**A**) Left: Mean Brier score per trial. Right: Mean accuracy score per trial. Dotted line indicates the parameter used for the final model (C = 1.99). (**B**) Corresponding between predicted cell types and reference cell types of 20% of the reference data after using 80% of the reference data for optimizing regularization strength in logistic regression. (**C**) Individual cells from training set (Red circles, 80% of the reference data), test set (Yellow circles, 20% of the reference data), and negative control (Black crosses, 20% of the reference data with random permutation of the selected gene set) plotted on a wheel plot. (**D**) Violin plot showing genes enriched in clusters from day 21 and 28 of differentiation. Log-library size normalized gene expression is shown.

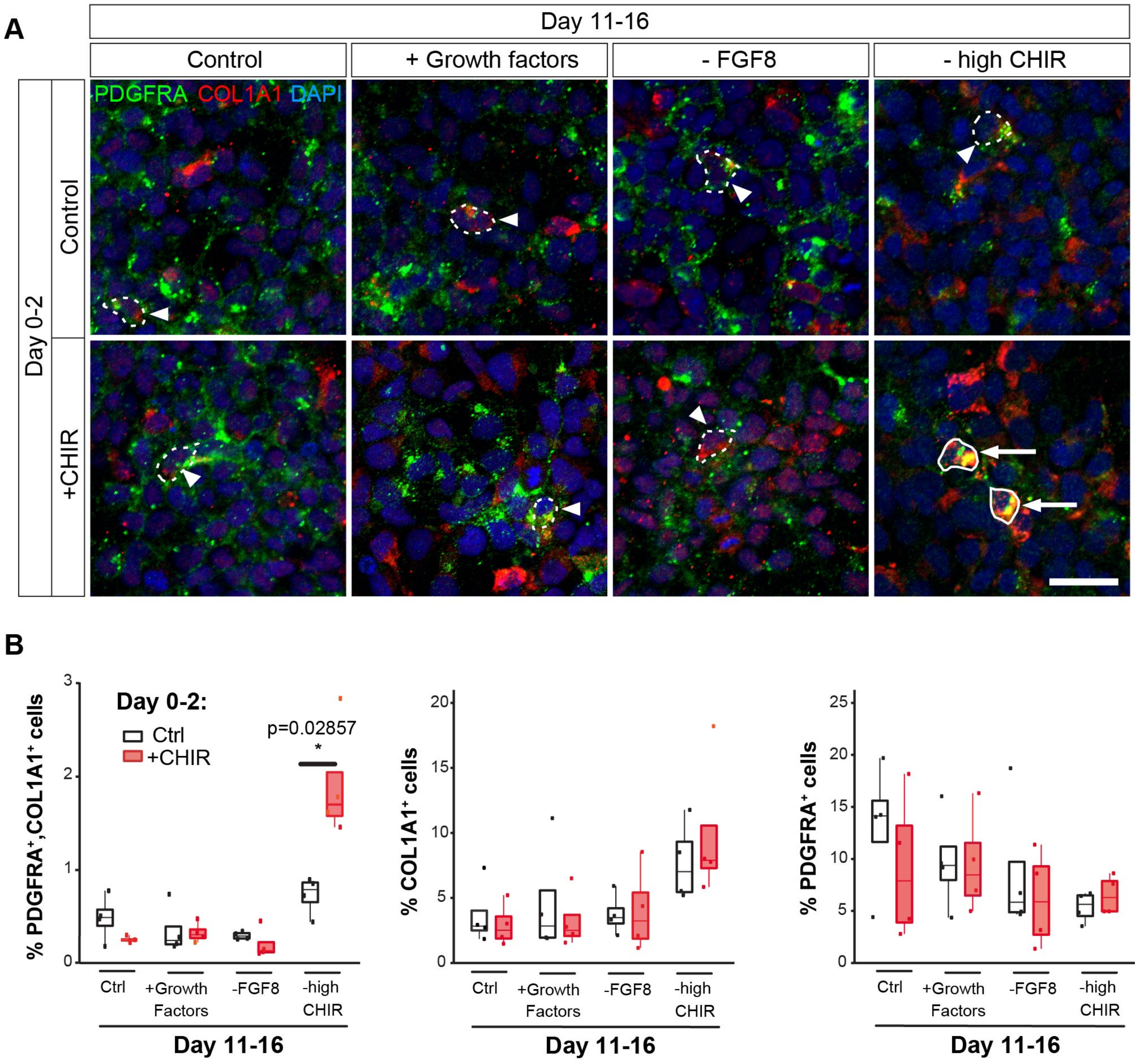


**Figure S7. VLMCs emerge after abnormal patterning with CHIR99021, related to Figure 5.** (**A**) Fluorescence immunocytochemistry staining for PDGFRA and COL1A1 showing that VLMCs, defined by the co-localization of PDGFRA and COL1A1, are not present in our standard culture conditions (control) or after adding growth factors or removing FGF8b at day 16. Note that only background levels of co-localization (few pixels) were found in few cells (dotted line and arrowhead). However, abnormal Wnt patterning with CHIR99021 too early (Day 0-2) and too short (until day 11) give rise to detectable VLMCs (encircled cells pointed by arrows). (**B**) Percentage of PDGFRA$^+$, COL1A1$^+$ and PDGFRA$^+$/COL1A1$^+$ cells out of the total cells (DAPI) in different conditions. Values under 1% PDGFRA$^+$/COL1A1$^+$ cells showed only background levels of staining (few pixels). Mann–Whitney U-test was used for pair comparison. Scale bar, 25 μm.

**Table S1**

**List of primer sequences**

| **Gene** | **Forward** | **Reverse** |
|---|---|---|
| *ABCA1* | ATGTGAGGCGGGAAAGACAGAG | ATCCTGTCAACAGCAGGCTTCC |
| *ALDH1A1* | TGTTAGCTGATGCCGACTTG | CTGGCCCTGGTGGTAGAATA |
| *BARHL1* | CCAGAACCGCAGGACTAAATGG | CTGGAGCGCTGAGTAATTGCCT |
| *CALB1* | GACGGAAGTGGTTACCTGGA | TGCCCATACTGATCCACAAA |
| *CORIN* | CATATCTCCATCGCCTCAGTTG | GGCAGGAGTCCATGACTGT |
| *DCX* | ACCTCCAGCAGCCAGCTCTCTA | GGCAGGTACAGGTCCTTGTGCT |
| *DEAF1* | *CCAGGTCCTCAGTCTCCTCCAA* | TGTCGTACACAGAAGGGTCCCA |
| *DKK3* | *GGTGGAAGAGATGGAGGCAGAA* | *CCAACCTTCGTGTCTGTGTTGG* |
| *EBF1* | GTGCGAGTTCATCGTCTGAGA | ACTTGTATCAGATTACTCTC |
| *EBF2* | GATTTGCTGGCAACGTTGGG | TCATTATTGGTCCATCAGAG |
| *EN1* | CGTGGCTTACTCCCCATTTA | TCTCGCTGTCTCTCCCTCTC |
| *ERBB4* | TGGCCACCAAACATGACTGACT | GAGAGGTGATGCCCTGTTGCTT |
| *FGF8B* | AGGTAACTGTTCAGTCCTCACC | TGTAGAGTTGGTAGGTCCGG |
| *FOXA2* | TTCAGGCCCGGCTAACTCT | AGTCTCGACCCCCACTTGCT |
| *FOXG1* | GCTGGACATGGGAGATAGGA | GTGGTGGTTGTCGTTCTGG |
| *GAPDH* | TTGAGGTCAATGAAGGGGTC | GAAGGTGAAGGTCGGAGTCA |
| *GBX2* | GTTCCCGCCGTCGCTGATGAT | GCCGGTGTAGACGAAATGGCCG |
| *KCNJ6* | TAGAGGACCCCTCCTGGACT | TCCCTCTGGGCATTTATCTG |
| *HOXA2* | AGTCTCGCCTTTAACCAGCA | TAGGCCAGCTCCACAGTTCT |
| *LMO3* | CTCTCAGTCCAGCCAGACACCA | GGCACACTTCAGGCAGTCTTCA |
| *LMX1A* | GATCCCTTCCGACAGGGTCTC | GGTTTCCCACTCTGGACTGC |
| *MSX1* | CGAGTTAAAGATGGGGAAACTG | GAGACATGGCCTCTAGCTCTGT |
| *NANOG* | ACAACTGGCCGAAGAATAGCA | GGTTCCCAGTCGGGTTCA |
| *NHLH1* | CCCGACAAGAAGCTCTCCAAGA | CAGGCTGAGTTCAGACGTCCAG |
| *NGN2* | GCTGGGTCTGGTACACGATT | GGCCTTCAGTCTACGGGTCT |
| *NKX2.1* | AGAGGGCTCTGTGCTGACAT | CAGAGTGTGCCCAGAGTGAA |
| *NEUROD1* | ACCCCTACTCCTACCAGTCGCC | GGCTTAACGTGGAAGACATGGG |
| *NR4A2* | CAGCTCCGATTTCTTAACTCCAG | GGTGAGGTCCATGCTAAACTTGA |
| *OTX2* | ACAAGTGGCCAATTCACTCC | GAGGTGGACAAGGGATCTGA |
| *PBX1* | TAAAAAGCCTTGGTGCTTCCCA | GCTCGTCCATCTCCAAAGGCTA |
| *PITX2* | CATGTCCACACGCGAAGAAATC | CCCGACGATTCTTGAACCAAAC |
| *POU5F1* | AGGGCCCCATTTTGGTACC | TCAGTTTGAATGCATGGGAGAGC |
| *POU6F1* | GCCTACAGCCAGTCAGCCATCT | GTTCCGCAGTTCAGCTTCGTTT |
| *SIX3* | AACTTCCGCGACCTCTACCACA | AGCTTCTCGGCCTCCTGGTAGT |
| *SOX2* | CAAGATGCACAACTCGGAGA | GCTTAGCCTCGTCGATGAAC |
| *SREBF1* | AACACAGACGTGCTCATGGAGG | CTCTGGAAAGGTGAGCCAGCAT |
| *TH* | ACTGGTTCACGGTGGAGTTC | TCTCAGGCTCCTCAGACAGG |
| *TUBB3* | CATTCTGGTGGACCTGGAAC | ATACTCCTCACGCACCTTGC |
| *WNT1* | GAGCCACGAGTTTGGATGTT | TGCAGGGAGAAAGGAGAGAA |
| *WNT5A* | ACTGCAAGTTCCACTGGTGCTG | GTGGCACCCACTACTTGCACAC |
| *WNT7A* | CAATCGGGACTATGAACCGGAA | GCCCAGAGCTACCACTGAGGAG |
| *WNT11* | GAAGCGACAGCTGCGACCTTAT | CAGGTGACGTAGCAGCACCAGT |
